# Phenotypic variation within and across transcriptomic cell types in mouse motor cortex

**DOI:** 10.1101/2020.02.03.929158

**Authors:** Federico Scala, Dmitry Kobak, Matteo Bernabucci, Yves Bernaerts, Cathryn René Cadwell, Jesus Ramon Castro, Leonard Hartmanis, Xiaolong Jiang, Sophie Laturnus, Elanine Miranda, Shalaka Mulherkar, Zheng Huan Tan, Zizhen Yao, Hongkui Zeng, Rickard Sandberg, Philipp Berens, Andreas Savas Tolias

**Affiliations:** Center for Neuroscience and Artificial Intelligence, Baylor College of Medicine, Houston, Texas, USA; Department of Neuroscience, Baylor College of Medicine, Houston, Texas, USA; Institute for Ophthalmic Research, University of Tübingen, Germany; International Max Planck Research School for Intelligent Systems, Germany; Department of Anatomic Pathology, University of California San Francisco, San Francisco, California, USA; Department of Cell and Molecular Biology, Karolinska Institutet, Stockholm, Sweden; Jan and Dan Ducan Neurological research Institute, Houston, TX, USA; Allen Institute for Brain Science, Seattle, Washington, USA; Center for Integrative Neuroscience, University of Tübingen, Germany; Institute for Bioinformatics and Medical Informatics, University of Tübingen, Germany; Bernstein Center for Computational Neuroscience, University of Tübingen, Germany

## Abstract

Cortical neurons exhibit astounding diversity in gene expression as well as in morphological and electrophysiological properties. Most existing neural taxonomies are based on either transcriptomic or morpho-electric criteria, as it has been technically challenging to study both aspects of neuronal diversity in the same set of cells. Here we used Patch-seq to combine patch-clamp recording, biocytin staining, and single-cell RNA sequencing of over 1300 neurons in adult mouse motor cortex, providing a comprehensive morpho-electric annotation of almost all transcriptomically defined neural cell types. We found that, although broad families of transcriptomic types (*Vip*, *Pvalb*, *Sst*, etc.) had distinct and essentially non-overlapping morpho-electric phenotypes, individual transcriptomic types within the same family were not well-separated in the morpho-electric space. Instead, there was a continuum of variability in morphology and electrophysiology, with neighbouring transcriptomic cell types showing similar morpho-electric features, often without clear boundaries between them. Our results suggest that neural types in the neocortex do not always form discrete entities. Instead, neurons follow a hierarchy consisting of distinct non-overlapping branches at the level of families, but can form continuous and correlated transcriptomic and morpho-electrical landscapes within families.

## Introduction

As animals can be grouped into species and assembled in a hierarchy of phylogenetic relationships to form the “tree of life”, neurons in the brain are thought to form discrete cell types, which in turn can be cast in a hierarchy of neuronal families and classes. The current view is that a neuronal cell type is characterized by a common genetic profile, giving rise to distinct physiological and anatomical properties including connectivity patterns (Masland, 2004; Zeng and Sanes, 2017). A comprehensive multi-modal taxonomy of neurons would resemble a parts list of the brain, helping us to decipher its bewildering complexity (Ecker et al., 2017; Mukamel and Ngai, 2019).

For more than a century, neurons have been classified into types by their anatomical and physiological characteristics, and more recently by molecular markers (Harris and Shepherd, 2015; Tremblay et al., 2016; Kepecs and Fishell, 2014; Rudy et al., 2011). In the last years, development of high-throughput single-cell sequencing techniques allowed to identify dozens of neural types based on their transcriptional profiles (Tasic et al., 2016, 2018; Zeisel et al., 2015, 2018), but linking the transcriptomically defined cell types to their phenotypes has remained a major challenge (Huang and Paul, 2019). At the same time, to understand the role that transcriptomic neural types play in cortical computations, it is necessary to know their anatomy, connectivity, and electrophysiology (Zeng and Sanes, 2017).

The BRAIN initiative cell census network (BICCN) aims at fully characterizing the cellular taxonomy of neurons in mouse motor cortex (MOp). In parallel work, Yao et al. perform an integrated transcriptomic analysis of the MOp, identifying 90 distinct neural types. To provide a comprehensive account of the anatomy and physiology of these transcriptomically defined cell types, we used the recently developed Patch-seq technique (Cadwell et al., 2016, 2017; Fuzik et al., 2016; Földy et al., 2016) and sampled over 1300 neurons from all cortical layers, combining single-cell RNA-sequencing, patch-clamp recordings and biocytin stainings in the same neurons.

Our data set covers all major families of excitatory and inhibitory cortical neurons and describes morpho-electric phenotypes for most of the transcriptomic cell types. Our analysis suggests that transcriptomic families have largely distinct phenotypes, but uncovers continuous morphoelectric variation within most major families.

## Results

### Patch-seq profiling of mouse motor cortex

We used Patch-seq (Cadwell et al., 2016, 2017) to profile neurons transriptomically, electrophysiologically and anatomically (Figure S1) across all layers of adult mouse MOp (mostly post-natal day P50+, median age P75, Figure S2a) using various Cre driver lines (Figure S3) to sample as diverse a population of neurons as possible. Neurons in acute slices were patch-clamped and stimulated with brief current impulses to record their electrophysiological activity, filled with biocytin for subsequent morphological recovery and reconstruction, and their RNA was extracted and sequenced using the Smart-Seq2 protocol (Picelli et al., 2013). In total, we performed whole-cell recordings from over 2000 cells, of which 1320 cells (from 262 mice) passed initial quality control and their mRNA was sequenced, yielding on average 1.3 million reads (median; mean ± SD on a log10 scale: 6.0 ± 0.6) and 6.8 ± 2.7 thousand (mean ± SD) detected genes per cell (Figure S2d). Of these, 642 neurons had sufficient staining and their morphologies were reconstructed.

Using the gene expression profiles, we mapped all sequenced neurons to the transcriptomic cell types (t-types) identified based on dissociated cells in a parallel work within the BICCN consortium (Yao et al., in preparation). To assign cell types, we used a nearest centroid classifier with Pearson correlation of log-expression across the most variable genes as a distance metric (Figure S4). Bootstrapping over genes was used to assess mapping confidence. The mapping was done separately using each of the seven reference data sets obtained with different sequencing technologies including single-cell and single-nucleus, Smart-seq2 and 10x (Yao et al., in preparation). We found that Patch-seq expression profiles were most similar to the single-nucleus Smart-seq2 data (Figure S2g,h). At the same time, there was good agreement between t-type assignments based on Smart-seq2 and 10x reference data (Figure S2i), so consensus t-type assignment over all seven reference data sets was used for all subsequent analysis. Cells that showed poor mapping (mostly due to low read count) or potential RNA contamination were excluded (Figure S2f), leaving 1221 neurons for further analysis (814 inhibitory, 407 excitatory; 368 and 267 with morphological reconstructions respectively).

The resulting data set covered 78 out of the 90 neural t-types (Figure 1a), with 75 t-types having at least one morphologically reconstructed neuron. The coverage was comparably good for CGE- and MGE-derived (caudal and medial ganglionic eminence) interneurons and for excitatory neurons. Within-type distributions of soma depths (Figure 1b) were in good agreement with prior literature (Tasic et al., 2018) and with the layer-specific nomenclature of excitatory t-types, confirming the validity of our t-type assignment. Positioning all cells on reference maps made with t-SNE (Maaten and Hinton, 2008; Kobak and Berens, 2019) also showed good overall coverage (Figure 1c–e, S5) with only few conspicuously uncovered regions (see Discussion below).

**Figure 1:**
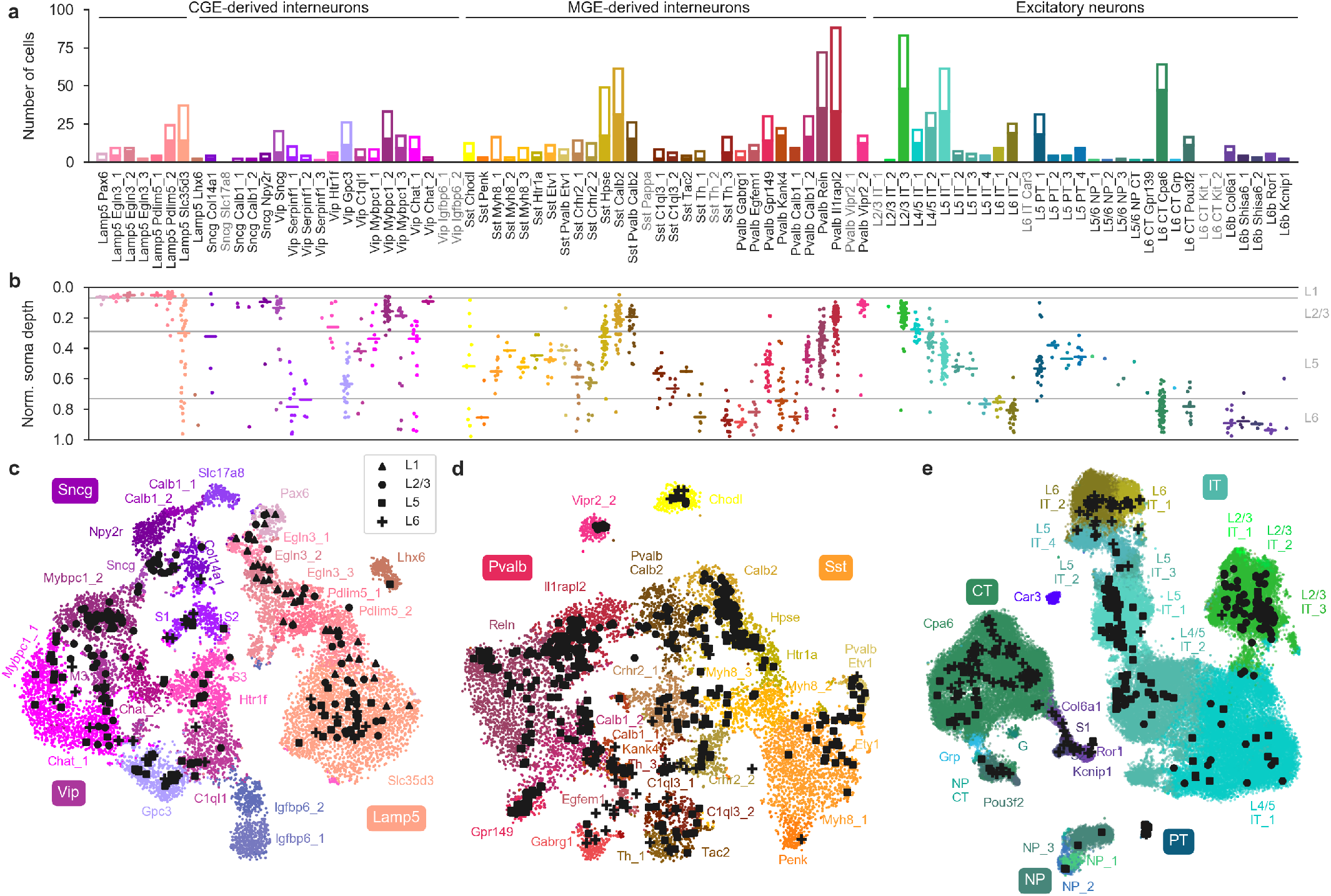
Transcriptomic coverage. **(a)** Number of Patch-seq cells assigned to each of the neural transcriptomic types (t-types) (Yao et al., in preparation). Colors are taken from the original publication, as well as the order of types. The filled part of each bar shows the number of morphologically reconstructed neurons. T-types with zero cells are shown with grey labels. Total number of neurons: 1221. **(b)** Normalized soma depths of all neurons of each t-type. For t-types with at least 3 cells, medians are indicated by horizontal lines. Soma depths were normalized by the cortical thickness in each slice (0: pia, 1: white matter). Grey horizontal lines indicate approximate layer boundaries identified via Nissl staining (L1: 0.07, L2/3: 0.29, L5: 0.73). Total number of neurons: 1181 (for some cells soma depth could not be measured due to failed staining). **(c)** T-SNE representation of CGE-derived interneurons from the single-cell 10x v2 reference dataset (*n* = 15 511; perplexity 30). T-type names are shortened by omitting the first word; some are abbreviated. Patch-seq cells from the *Vip*, *Sncg*, and *Lamp5* families were positioned on this t-SNE atlas (Kobak and Berens, 2019) and are shown as black symbols. Markers indicate layer, see legend. **(d)** Like (c), but for MGE-derived interneurons (*n* = 12 083; perplexity 30). **(e)** Like (c), but for excitatory neurons (*n* = 93 829; perplexity 100). A single t-SNE embedding with all cells from panels (c–e) is shown in Figure S5.

The observed phenotypes included most of the morphological and electrophysiological types of cortical neurons previously described in mice and rats (Jiang et al., 2015; Markram et al., 2015; Gouwens et al., 2019), allowing us to link transcriptomic and morpho-electric descriptions of the neural landscape (Figure 2; see Supplementary File 1 for all reconstructions and Figure S6 for interneurons mapped to the t-type taxonomy from Tasic et al. (2018)). To obtain quantitative characterizations of the morpho-electric phenotypes, we automatically extracted 28 electrophysiological (Figure S7) and ~100 morphological features for each cell. In the next section, we provide a detailed description of all t-types with sufficient coverage.

**Figure 2:**
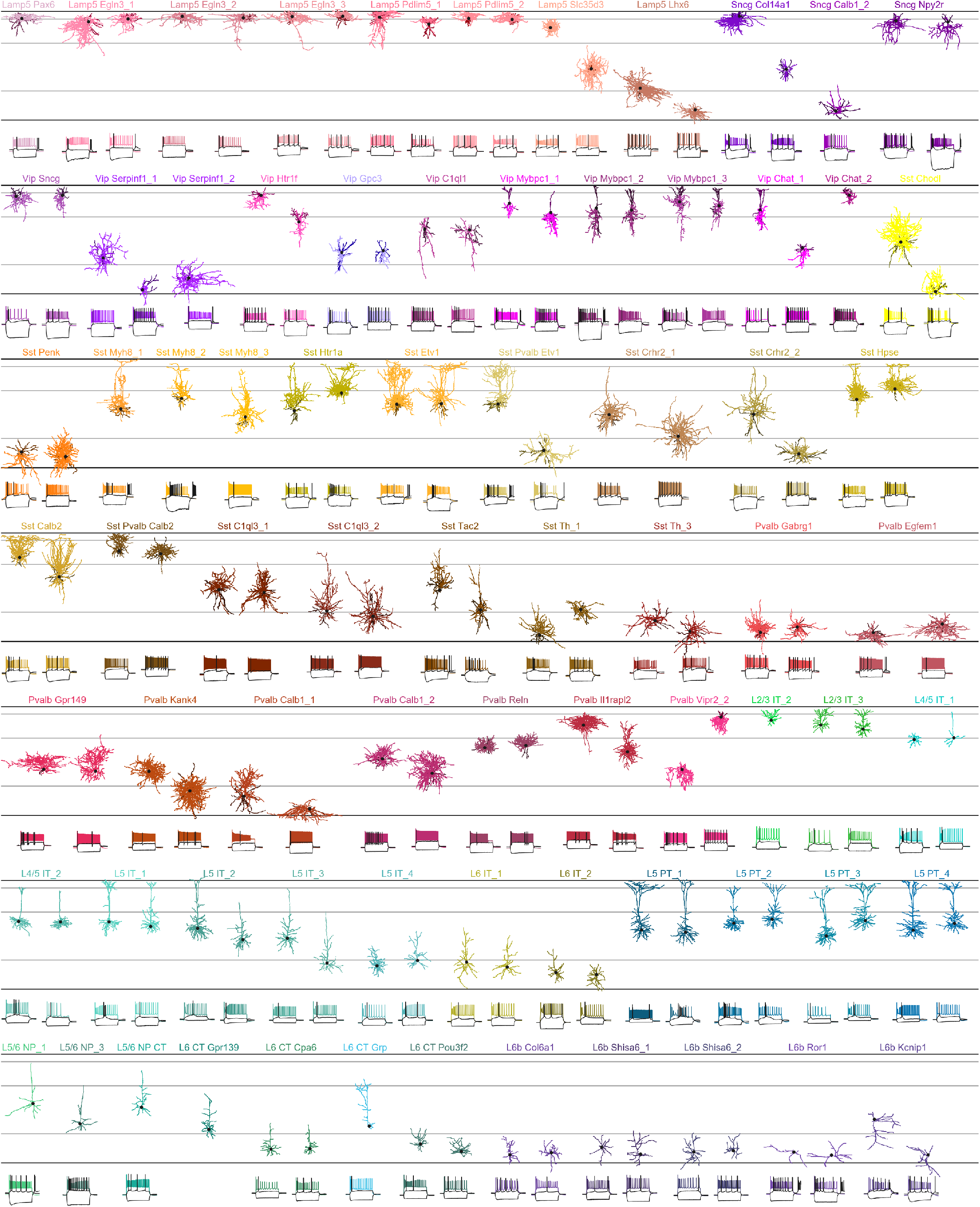
Diversity of mouse cortical neurons. Two representative examples per t-type, or one if only one reconstruction was available. In total 137 neurons in 75 t-types. For interneurons, dendrites are shown in darker colors. For excitatory neurons, only dendrites are shown. Black dots mark soma locations. Horizontal grey lines show approximate layer boundaries. Three voltage traces are shown for each neuron: the hyperpolarization trace obtained with the smallest current stimulation, the first depolarization trace eliciting at least one action potential, and the depolarization trace showing maximal firing rate. Stimulation length: 600 ms. The length of the shown voltage traces: 900 ms. Electrophysiological recording for one neuron did not pass quality control and is not shown.

### Morpho-electric phenotypes of transcriptomically defined neuron types

#### CGE-derived interneuron types

The *Lamp5* family mostly consisted of L1 interneurons. The *Lamp5 Egln3_1* and *Lamp5 Egln3_2* types likely corresponded to previously described alpha7 and canopy cells respectively (Schuman et al., 2019) based on the expression of known marker genes (*Ndnf^−^ Chrna7*^+^ and *Ndnf*^+^ *Npy*^−^; here and below reported marker gene expression is based on the data from Yao et el.) and electrophysiology: *Lamp5 Egln3_1* was characterized by larger membrane time constant and hyperpolarization sag, stronger bursts, and rebound firing. The *Lamp5 Pdlim 5_2* type (*Ndnf*^+^ *Npy*^+^) corresponded to late-spiking neurogliaform cells (NGCs) in L1 with wide asymmetric action potentials (APs) and deep afterhyperpolarization (AHP). NGCs in L2/3, L5, and L6 belonged to the *Lamp5 Slc35d3* type (*Ndnf*^−^) that showed layer-adapting axonal morphology. The transcriptomically isolated *Lamp5 Lhx6* type (cf. Figure 1c) consisted of deep L5/L6 neurogliaform-like cells with NGC morphology and deep AHP but narrow APs. As the *Lamp5 Lhx6* type is putatively MGE-derived (Tasic et al., 2018), this suggests that although all deep NGCs belong to the *Lamp5* family, some are CGE- and some are MGE-derived [as is the case in hippocampus (Tricoire et al., 2010; Pelkey et al., 2017; Harris et al., 2018)], resolving an open question in the field (Huang and Paul, 2019).

The *Sncg* family (mostly *Vip*^−^ and strongly *Cck*^+^) proved difficult to sample due to the lack of specialized Cre lines; we only obtained 10 cells with 5 reconstructions. We found them in all layers from L1 to L6 with diverse morphologies; they mostly showed irregular bursting firing, sometimes with a strong rebound. Several cells in the upper L2/3 had large axonal morphologies, likely corresponding to ‘large *Cck* basket cells’ (Tremblay et al., 2016).

The *Vip* family was most abundant in L2/3, in agreement with the literature (Prönneke et al., 2015). Most L2/3 neurons from this family belonged to the *Vip Sncg*, *Vip Mybpc1_2*, and *Vip Mybpc1_3* types. *Vip Sncg* (strong *Cck*^+^ expression) neurons had local dendritic and axonal morphologies, identifying them as ‘small *Cck* basket cells’ (Tremblay et al., 2016). The other two types tended to have more vertically oriented morphologies, sometimes with bipolar dendritic structure, and showed diverse firing patterns with some neurons exhibiting large membrane time constant, hyperpolarization sag and strong rebound firing. All three types were also characterized by high input resistance (Tremblay et al., 2016).

In L5 we found *Vip Mybpc1_1* (*Calb2*^+^), *Vip Chat_1* (*Calb2*^+^ *Chat*^+^), *Vip Gpc3*, *Vip Htr1f* and *Vip C1ql1* types. Their axons and dendrites remained mostly local (Prönneke et al., 2015), with some cells having deep-projecting axons (Jiang et al., 2015; Prönneke et al., 2015). *Vip Mybpc1_1* and *Vip Chat_1* also contained some bipolar cells in L2/3 and upper L5. Despite *Vip* neurons traditionally being characterized by their high input resistance (Tremblay et al., 2016), some of these L5 types, especially *Vip Gpc3*, showed only moderate input resistance values, comparable to the *Sst* family. This type also had particularly low resting membrane potential. The three *Vip Serpinf1* types (*Cck*^+^) were found in deep L5 and L6. We did not obtain any cells from the *Vip Igfbp6* types, presumably due to their very weak *Vip* expression (Tasic et al., 2018).

#### MGE-derived interneuron types

The *Sst* family in L2/3 was mostly represented by the *Sst Calb2* type, with Martinotti morphology and adapting firing pattern (Tremblay et al., 2016; Urban-Ciecko and Barth, 2016). In upper L5, the cells of this type typically showed ‘fanning-out’ Martinotti morphology (Muñoz et al., 2017; Nigro et al., 2018). The neighbouring *Sst Hpse* type also showed ascending axons typical of Martinotti cells but with denser local axons and sparser ‘fanning-out’ projections to L1. Both types were distinguished from other *Sst* t-types by non-zero afterdepolarization (ADP). Between the two, *Sst Hpse* had smaller membrane time constant, in agreement with earlier findings in visual cortex (Scala et al., 2019). *Sst Htr1a* appeared very similar to its neighboring *Sst Hpse*. In addition, we found *Sst Pvalb Calb2* neurons in L2/3. This t-type is transcriptomically in between the *Sst* and the *Pvalb* families, and we found it to be in between also in terms of the morpho-electric phenotype (see also below). These neurons showed lower AP width and higher firing rate than typical for the *Sst* family (Scala et al., 2019). Furthermore, some cells in this type had Martinotti morphology while some others looked like typical L2/3 basket cells.

Multiple *Sst* types were predominantly found in L5, showing diverse firing patterns and morphologies. *Sst Myh8_1*, *Sst Etv1*, and *Sst Pvalb Etv1* showed ‘T-shaped’ Martinotti morphologies (Muñoz et al., 2017; Nigro et al., 2018) and strong rebound firing. The *Sst Pvalb Etv1*, together with *Sst Myh8_3* exhibited strong hyperpolarization sag. The two *Sst C1ql3* types showed non-Martinotti morphology without ascending axons (Tremblay et al., 2016; Gouwens et al., 2019) and deep AHP. The two *Sst Crhr2* types showed mostly local axonal arbor but with some sparse ascending axons.

In L6, the *Sst* family was represented by the *Sst Th* types and the *Sst Penk* type. All of them had mostly local axonal arborization within L6 (Perrenoud et al., 2012). Finally, we found cells of the transcriptomically isolated *Sst Chodl* type in all layers from upper L2/3 down to the bottom of L6. This type is thought to have long-range projections (Tasic et al., 2018; Gouwens et al., 2019) and for two cells in L6 we could indeed see an axon disappearing into the white matter. The neurons of this type had low rebound potential and low hyperpolarization sag but large variability in the membrane time constant.

The *Pvalb* family is known to consist of fast-spiking (FS) chandelier cells targeting the axons of excitatory neurons, and of soma-targeting FS basket cells (Tremblay et al., 2016). The chandelier cells have an easily recognizable axonal morphology with straight terminal ‘cartridges’ (Tremblay et al., 2016). We found chandelier cells exclusively in the transcriptomically isolated *Pvalb Vipr2_2* type, confirming its previously postulated identity (Tasic et al., 2018). They were mostly located in upper L2/3 close to the L1 boundary, but deep L5 chandelier cells belonged to the same t-type. In terms of electrophysiology, chandelier cells had a lower firing rate compared to the basket cells, and a practically absent hyperpolarization sag.

The rest of the *Pvalb* family consisted of FS basket cells with various axonal morphologies and layer distributions (Jiang et al., 2015; Scala et al., 2019; Gouwens et al., 2019). In L2/3, most cells were of *Pvalb Il1rapl2* type, with classical L2/3 basket morphology (Jiang et al., 2015). We did not encounter double-bouquet basket cells, previously described in L2/3 of mouse V1 (Jiang et al., 2015). In L5, the same work distinguished between large basket cells, small (or shrub) basket cells, and horizontally elongated basket cells, with differing connectivity. We found them preferentially in *Pvalb Il1rapl2*, *Pvalb Reln*, and *Pvalb Gpr149* types respectively, but with substantial overlap (see below). Notably, *Pvalb Il1rapl2* type showed strongly layer-adapting morphologies, with L2/3 neurons looking very different from L5 (Supplementary File 1). *Pvalb Calb1_2* and *Pvalb Kank4* showed a variety of axonal morphologies, including some with large local arborization with dense spherical shape. In L6, FS cells belonged to the *Pvalb Gabrg1*, *Pvalb Egfem1*, *Pvalb Calb1_1* and *Pvalb Kank4*, and mostly had local axons (Perrenoud et al., 2012) [we did not encounter L6 FS cells with translaminar axons reaching up to L1, that were reported in V1 (Bortone et al., 2014; Gouwens et al., 2019)]. Some of the deep neurons exhibited a horizontally elongated or downward projecting axon mostly innervating L6b (Gouwens et al., 2019). *Pvalb Gabrg1* and *Pvalb Egfem1* types were characterized by larger hyperpolarization sag and rebound potential compared to the other FS neurons.

#### Excitatory neuron types

Transcriptomically, cortical excitatory neurons are classified into the well-separated intertelencephalic (IT), pyramidal tract (PT, also called ET), corticothalamic (CT), and near-projecting (NP) families (Tasic et al., 2018). Morphologically, they have been classified into big-tufted, small-tufted, untufted neurons, depending on the shape of the apical dendrite tuft, stellate neurons without an apical dendrite, and horizontal/inverted neurons in L6 (Oberlaender et al., 2011; Marx and Feldmeyer, 2012; Wang et al., 2018; Kanari et al., 2019).

The IT neurons in L2/3 were tufted pyramidal cells with high rheobase and almost all of them were assigned to the *L2/3 IT_3* type. The *L4/5 IT_1* cells were located on the boundary between L2/3 and L5 and possibly corresponded to the quasi-L4 neurons described previously in motor cortex (Yamawaki et al., 2014). Neurons of this type had diverse morphologies with some pyramidal and some stellate cells. The *L4/5 IT_2* cells mostly had a thin untufted apical dendrite, while *L5 IT_1* and *L5 IT_2* types contained deeper and larger tufted pyramidal neurons. L6 IT neurons were short and untufted; *L6 IT_1* showed pyramidal morphologies while *L6 IT_2* were often stellate or inverted (Zhang and Deschênes, 1997). The first type had broader APs than the second.

L5 PT neurons tended to be large big-tufted cells with the apical dendrite often bifurcating close to the soma, suggesting that these were corticospinal cells (Oswald et al., 2013; Ramaswamy and Markram, 2015). They had bigger hyperpolarization sag and higher rebound potential, compared to the L5 IT neurons. We did not observe consistent morpho-electric differences between the four PT types, but prior research suggests that they can have different projection targets (Economo et al., 2018; Tasic et al., 2018).

NP neurons proved very difficult to obtain without a specialized Cre driver line. The few neurons in our data set were all untufted, with sparse basal dendrites, in agreement with prior literature (Gouwens et al., 2019).

All CT t-types were preferentially located in L6 and had mostly untufted apical dendrites (Zhang and Deschênes, 1997). In line with previous literature (Thomson, 2010), they could be distinguished from L6 IT neurons by a lower inter-spike interval adaptation index (Figure S7). L6b types, transcriptomically related to the CT family (Figure 1e), were all stellate, inverted, or horizontal, located preferentially in the bottom of L6. The *L6b Ror1* type stood out, having horizontal dendritic morphology and showing strong rebound firing.

### Transcriptomic families have distinct morpho-electric phenotypes

We next asked to what extent electrophysiological phenotype could be predicted by gene expression across the entire data set. We focused on 16 well-behaved electrophysiological features and used sparse reduced-rank regression (Kobak et al., 2019), a technique that predicts the firing properties based on a low-dimensional latent space representation computed from a sparse selection of genes. We used cross-validation to tune the regularization strength (Figure S8). The selected model used 25 genes with a 5-dimensional latent space and achieved a cross-validated *R*^2^ of 0.40. To visualize the structure of the latent space, one can project gene expression and electrophysiological properties onto the latent dimensions (Figure 3). The cross-validated correlations between the first three pairs of projections were 0.90, 0.74, and 0.65 respectively.

**Figure 3:**
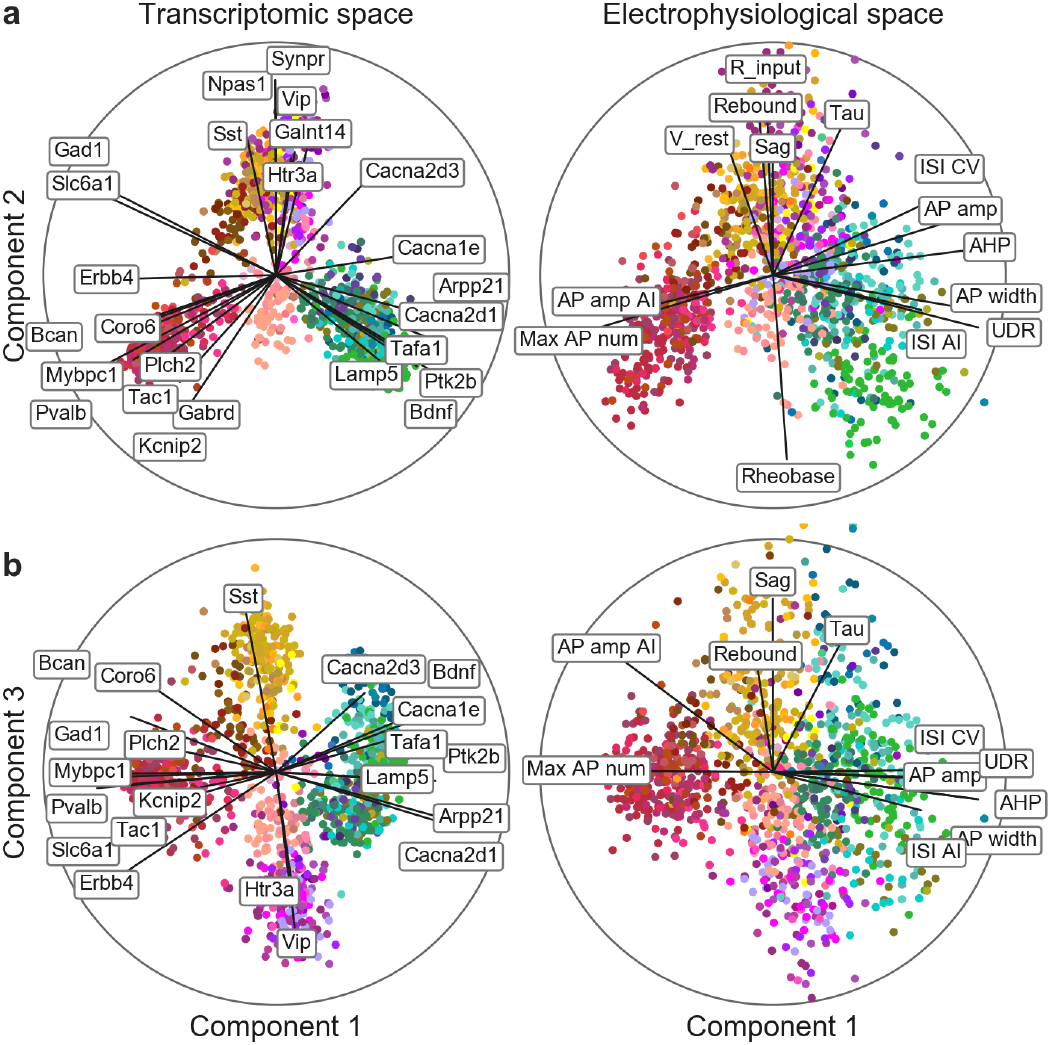
Sparse reduced-rank regression. A sparse RRR model (Kobak et al., 2019) to predict combined electrophysiological features from the gene expression. Transcriptomic data are linearly projected to a low-dimensional space that allows to reconstruct electrophysiological data; here shown are components 1 and 2 (a), and 1 and 3 (b), of rank-5 model. Sample size *n* = 1213. Color corresponds to the t-type. The model selected 25 genes, shown on the left. Each panel is a biplot, showing correlations of original features with both components; the circle corresponds to correlation 1. Only features with average correlation above 0.4 are shown. Abbreviations: ISI – interspike interval, CV – coefficient of variation, UDR – upstroke-to-downstroke ratio, AI – adaptation index, AP – action potential.

These first three components clearly separated five major neuron groups: the *Pvalb*, *Sst*, *Vip*, and *Lamp5* families, and the excitatory neuron class (Figure 3). These groups had distinct electrophysiological properties: for example, as expected, *Pvalb* neurons were characterized by high firing rates while *Sst* neurons had high values of the hyperpolarization sag and rebound (Figure 3, right). Some of the genes selected by the model were prominent marker genes, such as the pan-inhibitory markers *Gad1* and *Slc6a1* related to the gamma-aminobutyric acid (GABA) processing, or more specific inhibitory markers *Sst*, *Vip*, *Pvalb*, *Tac1*, or *Htr3a*. Interestingly, some other selected genes were more directly related to electrophysiological properties, such as calcium channel subunits *Cacna2d1* and *Cacna2d3* or potassium channel-interacting protein *Kcnip2*, which may modulate firing properties in individual families. A reduced-rank regression model restricted to using only ion channel genes (Figure S9) performed not much worse compared to the full model (cross-validated *R*^2^ = 0.35 and correlations 0.86, 0.70, and 0.55, with regularization set to select 25 genes).

Similarly, a 2D t-SNE embedding of Patch-seq cells based on the same electrophysiological features showed that major transcriptomic families have distinct electrophysiological properties (Figure 4a): the *Pvalb*, *Lamp5*, *Sst*, *Vip*, CT, IT, and PT families were mostly well-separated from each other, although there were no truly isolated clusters. We quantified this separation using a confusion matrix for *k*-nearest neighbors (kNN) classification of cells into families: it was mostly diagonal, with only the PT family strongly overlapping with the IT (Figure 4d).

**Figure 4:**
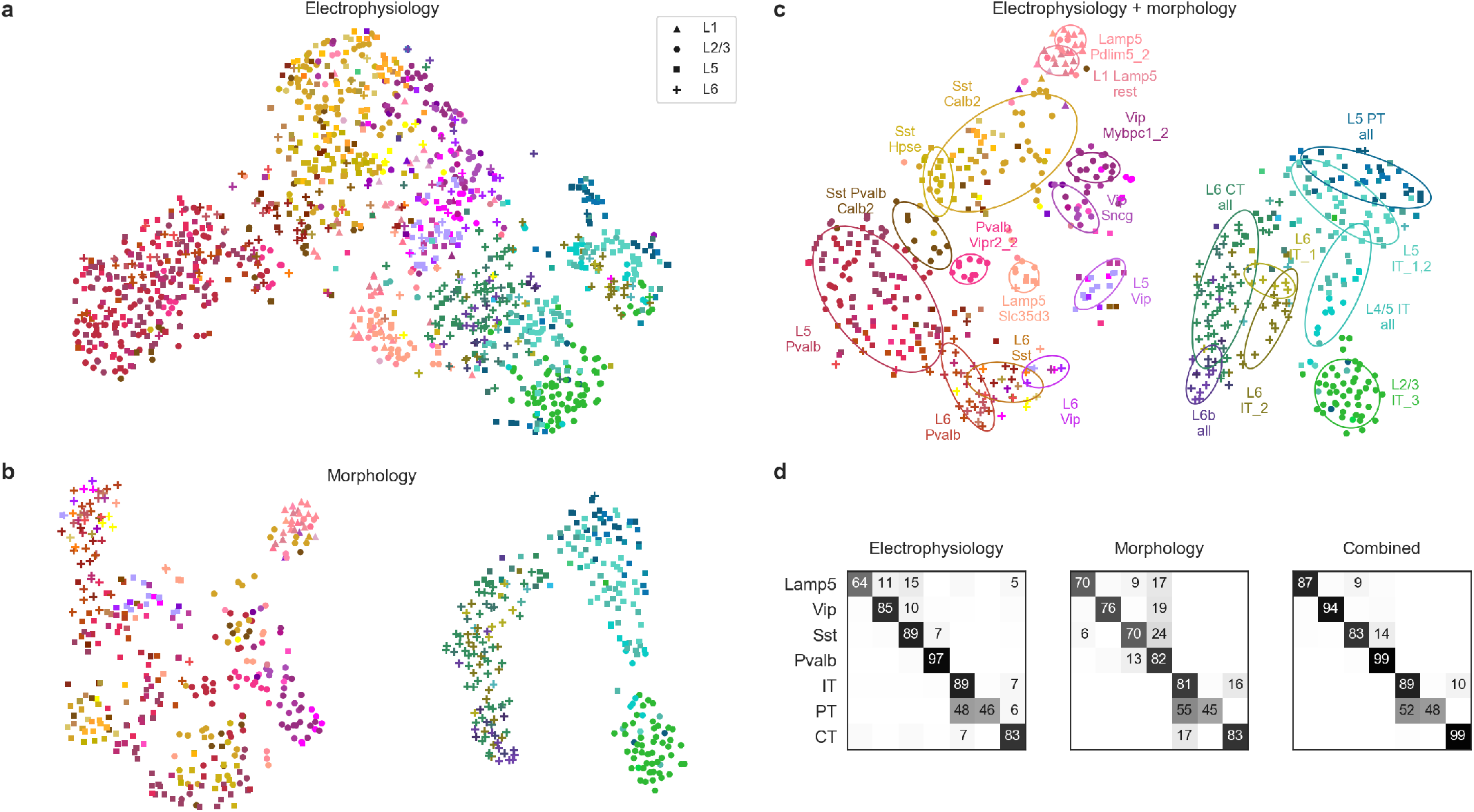
Morpho-electric t-SNE embeddings. **(a)** T-SNE embedding constructed using electrophysiological features. Color corresponds to the t-type, marker shape corresponds to the cortical layer (see legend). 1311 cells used to construct the embedding, 1213 cells with t-type labels shown. Perplexity 30. **(b)** T-SNE embedding constructed using combined morphometric features and *z*-profiles. Sample size: 633 cells. Perplexity 30. **(c)** T-SNE embedding constructed using combined electrophysiological and morphological features. Sample size: 625 cells. Perplexity 30. Ellipses show 80% coverage ellipses for the most prominent t-types, as well as for some groups of related t-types and some layer-restricted families. We chose these groups in order to reduce the overlap between ellipses. **(d)** Confusion matrices for classifying cells into seven transcriptomic families using kNN classifier (*k* = 10) and three different feature sets. Each row shows what fraction of cells from a given family gets classified in each of the seven families. The values in each row sum to 100%. Only values above 5% are shown.

We also constructed a 2D t-SNE embedding based on the morphological features (Figure 4b). We used only dendritic features for the excitatory cells, but both axonal and dendritic features for the inhibitory cells, leading to a strong separation between these two major classes. Within each class, cells were strongly segregated by the soma depth, with excitatory cells forming a mostly onedimensional manifold. The separability between inhibitory families was weaker than with electrophysiological features (Figure 4d). The between-family separability was the strongest when we combined electrophysiological and morphological features into a joint representation (Figure 4c,d), showing that these sets of properties are not redundant.

Together, these analyses suggest that different transcriptomic families had distinct morpho-electric phenotypes, in agreement with them being well-separated in the transcriptomic space (Tasic et al., 2018).

### Continuous phenotypic variation within transcriptomic families

Within individual transcriptomic families, morpho-electric phenotypes rarely formed isolated clusters (Figure 4). Moreover, we often found that morpho-electric phenotypes varied continuously from one t-type to another (Figure 5). For example, electrophysiological properties of the t-types within the *Vip* family varied continuously across the transcriptomic landscape, with e.g. the membrane time constant having its largest values close to the *Sncg* family and gradually decreasing all the way to the L5 area near *Vip Gpc3* (Figure 5a). For each pair of t-types we computed the transcriptomic distance (Euclidean distance between average log-counts in the reference data) and the electrophysiological distance (Euclidean distance between average feature vectors), and found that these two measures were correlated with *r* = 0.52 (Figure 5a, inset). This correlation was the highest for the input resistance (0.74) and the membrane time constant (0.61).

**Figure 5:**
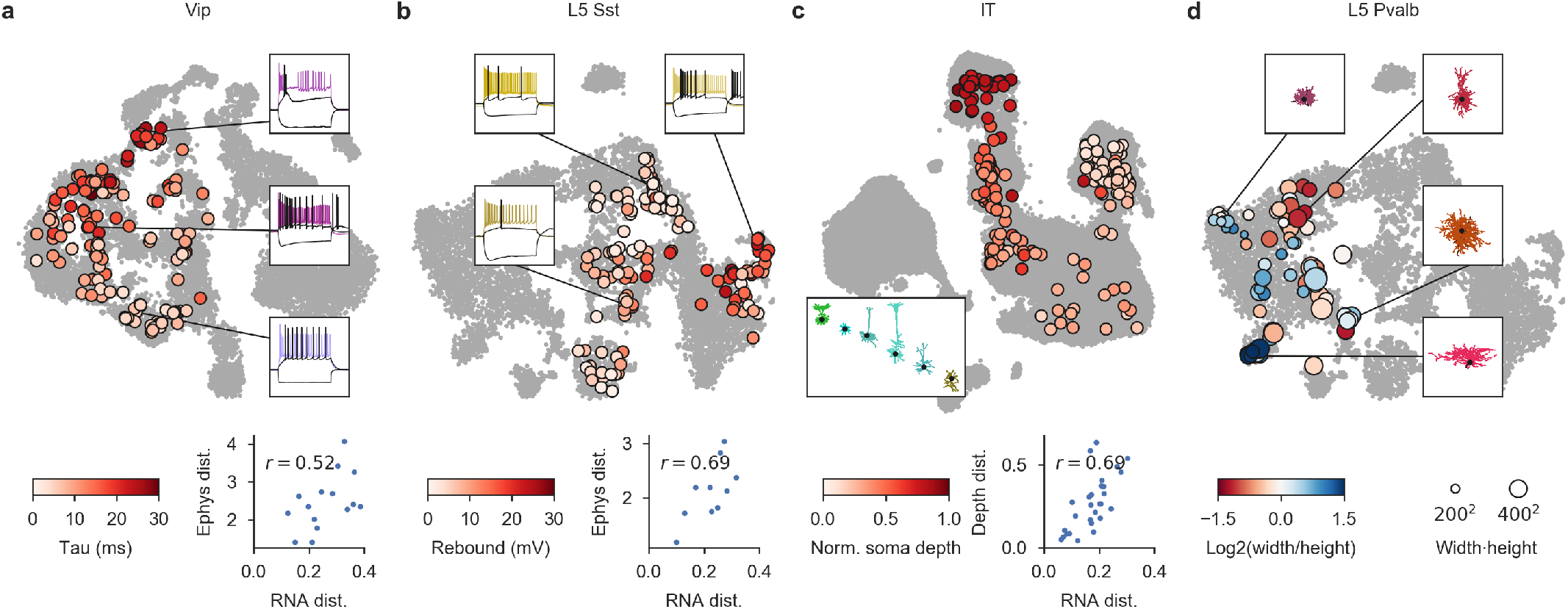
Phenotypic variability within transcriptomic families. **(a)** The *Vip* family neurons mapped to the reference t-SNE embedding as in Figure 1, colored by the membrane time constant. Insets show example firing traces. Scatter plot shows correlation between the transcriptomic distances and electrophysiological distances across all pairs of t-types (for 6 t-types with at least 10 cells). Transcriptomic distance was computed using the reference 10x data as the correlation between average log-expression across most variable genes. Electrophysiological distance is Euclidean distance between the average feature vectors. **(b)** The *Sst* family neurons from layer 5 (excluding *Sst Chodl* t-type), colored by the rebound. Scatter plot was done using 5 types with at least 10 cells. **(c)** IT neurons, colored by the normalized soma depth. Inset shows examples of IT neurons at different depths, colored by the t-type. Scatter plot was done using 8 t-types with at least 5 cells and shows correlation between the transcriptomic distances and the cortical depth distances. Cortical depth distance is Euclidean distance between the average normalized soma depths. **(d)** The *Pvalb* family neurons from layer 5, colored by the axonal width/height log-ratio. Circle area corresponds to the width/height product. Insets show some example morphologies.

The *Sst* family is known to be transcriptomically (Tasic et al., 2018) and phenotypically (Muñoz et al., 2017; Nigro et al., 2018; Naka et al., 2019) diverse in L5. Here we also found that electrophysiological properties varied continuously across the transcriptomic landscape, with neighbouring t-types consistently showing similar rebound values (Figure 5b). The transcriptomic and electrophysiological between-type distances were correlated with *r* = 0.69 (Figure 5c, inset), and among the individual features the correlation was the highest for the rebound (0.85), the AP amplitude (0.69), and the rheobase (0.65). Pooling all families together, transcriptomic and electrophysiological between-type within-family distances (*n* = 77 pairs) were highly correlated with *r* = 0.70 (Figure S10).

The IT family provides an example of a similar phenomenon in another data modality (Figure 5c). IT neurons span all layers from L2/3 to L6 and it is known that IT t-types are largely layer-restricted (Tasic et al., 2018). We found that L4/5 and L5 IT t-types that were transcriptomically close to the L2/3 IT t-types were located at the top of L5 close to the border between L2/3 and L5, whereas L5 IT t-types that were transcriptomically close to L6 IT t-types were located at the bottom of L5 close to the border with L6. Across the IT t-types, normalized soma depth varied smoothly with the transcriptome (*r* = 0.69; Figure 5c).

The *Pvalb* family is usually understood as electrophysiologically homogenous (all neurons are FS) but has been described as morphologically diverse, in particular in L5 (Jiang et al., 2015). However, it was previously unclear whether different morphologies such as shrub-like or horizontally elongated corresponded to different t-types. While we found that different t-types had different preferred morphologies (see above), they showed substantial overlap, in agreement with the L5 *Pvalb* t-types themselves not having clear boundaries (Tasic et al., 2018) (Figure 1d). The shape of the axonal arbor showed continuous changes across the transcriptomic landscape (Figure 5d): small shrub-like basket cells, horizontally elongated basket cells, and vertically elongated classical basket cells were located in different corners of the t-SNE embedding, with intermediate morphologies in between.

In summary, we found that within the major transcriptomic families, morpho-electric phenotypes and/or soma depth often varied smoothly between neighbouring t-types, indicating that transcriptomic neighbourhood relationships in many cases corresponded to similarities in other modalities.

### Individual t-types can have variable morpho-electric phenotypes

To study morpho-electric phenotypes of individual t-types, we measured (a) how consistently they conformed to their respective transcriptomic families (Figure 6a) and (b) how variable they were within a t-type (Figure 6b). First, we used a kNN classifier to classify cells from each t-type with at least 10 cells into transcriptomic families (the same as in Figure 4d). For this analysis we used electrophysiological, morphological, and combined features. When using morphological or combined features, we layer-restricted all t-types, to ignore the high between-layer morphological variability (see above).

**Figure 6:**
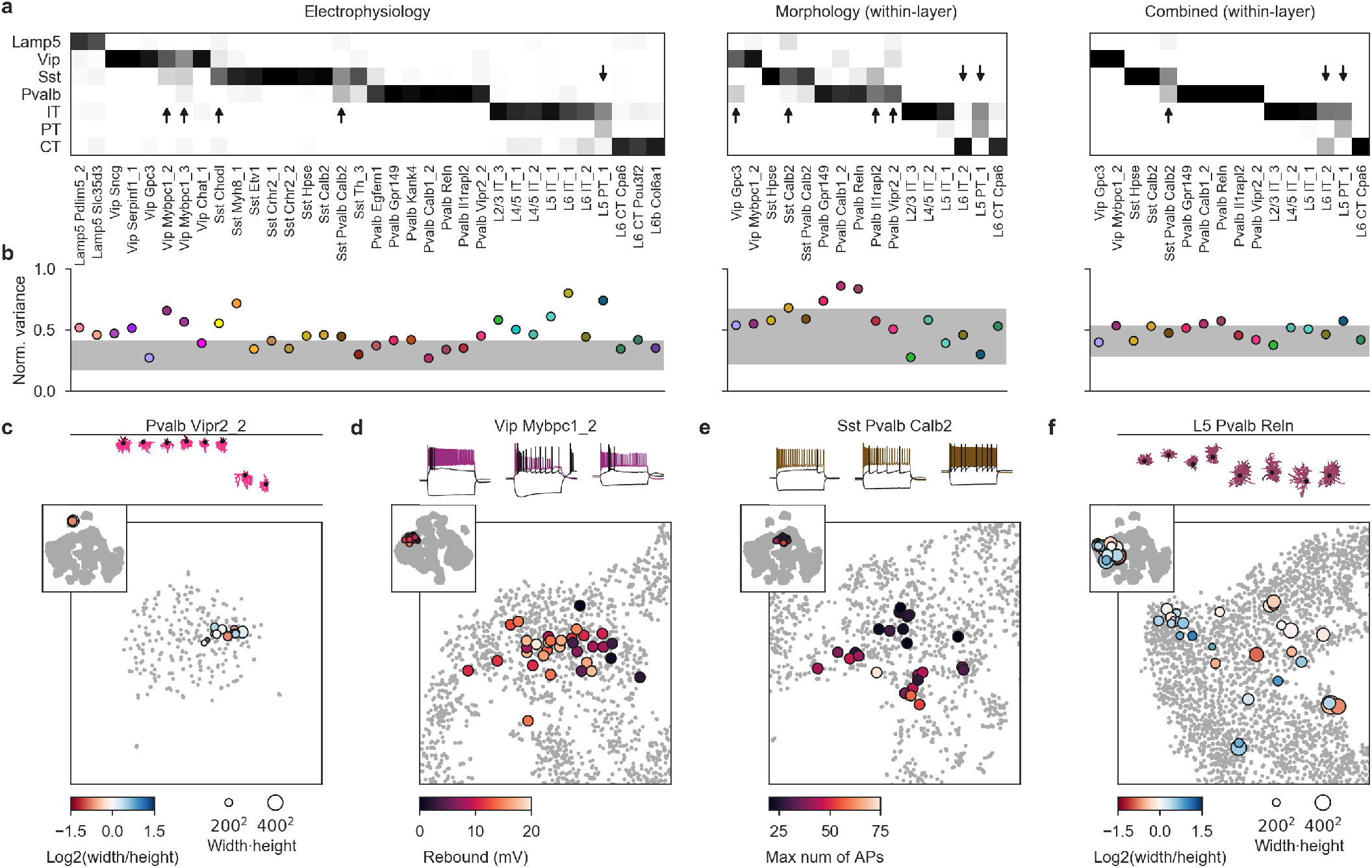
Phenotypic variability of individual t-types. **(a)** Confusion matrices for classifying cells from each t-type into seven transcriptomic families, using electrophysiological, morphological, and combined features. Only t-types with at least 10 cells are shown. For morphological and combined features we only took cells from one cortical layer. Values in each column map to 1. Arrows mark t-types that are classified into wrong families more often than 25% of the time. We used kNN-based classifier with *k* = 10. **(b)** Normalized total variance of features in each t-type. Higher values correspond to t-types with more variable phenotypes. Horizontal grey band shows the min/max normalized variances of *k*-means clusters. **(c)** Exemplary morphologies of *Pvalb Vipr2_2* chandelier neurons and t-SNE overlay colored by the axonal width/height log-ratio as in Figure 5d. **(d)** Three exemplary traces of cells from the *Vip Mybpc1_2* type (all with confidence above 90%) and t-SNE overlay colored by the rebound. **(e)** Three exemplary traces of cells from the *Sst Pvalb Calb2* (confidence above 90%) and t-SNE overlay colored by the maximum firing rate. **(f)** Exemplary morphologies of L5 cells from the *Pvalb Reln* type and t-SNE overlay colored by the axonal width/height log-ratio as in Figure 5d.

Most t-types could be unambiguously placed into the correct family (Figure 6a), but some t-types were in between two families with respect to their electrophysiological or morphological features. For example, many *Sst Pvalb Calb2* neurons were classified as belonging to the *Pvalb* family based on electrophysiology. Similarly, *Vip Mybpc1* neurons often had *Sst*-like firing, while *L6 IT_2* neurons had CT-like dendritic morphology. Thus, while overall, transcriptomic family was highly predictive of the cell phenotype, there were some t-types exhibiting morphological or physiological phenotypes reminiscent of another transcriptomic family.

Next, we measured the normalized total variance of each t-type using electrophysiological, morphological, or combined features and compared it to the normalized total variance of phenotype clusters derived by *k*-means clustering (with *k* set to the number of t-types). The rationale here was that the variance of the morpho-electric *k*-means clusters would reflect the minimal possible variance obtainable in our dataset. Values much above the cluster variances indicate non-trivial phenotypic variability within a t-type.

We found that many t-types had total variance substantially above the variances of the *k*-means clusters (Figure 6b) and an alternative analysis using entropies of Leiden clustering (Traag et al., 2019) often highlighted the same t-types as variable (Figure S11). Not all t-types showed high variability: some of them, such as the transcriptomically isolated *Pvalb Vipr2_2* type (chandelier cells), appeared morpho-electrically homogenous (Figure 6c). In contrast, *Vip Mybpc1_2* was marked as having high electrophysiological variability and indeed had high variance in input resistance, membrane time constant, and rebound (Figure S7). Overlaying the rebound values on the t-SNE embedding (Figure 6d) suggested that cells with low rebound were located close to the boundary with the *Vip Sncg* type that on average had low rebound, suggesting that in some cases, high within-t-type morpho-electric variability could be partially explained by the within-t-type transcriptomic variability. Similarly, *Sst Pvalb Calb2* cells had high variability in terms of the maximum firing rate, but high-firing cells were mostly grouped in one part of the transcriptomic landscape (Figure 6e).

Some t-types also showed pronounced within-t-type (and within-layer) morphological variability. For example, L5 cells from *Pvalb Reln* type exhibited a wide variety of different axonal morphologies (Figure 6f). At the same time, most of the small shrub-like cells were located in the upper-left corner of the t-SNE embedding, and those were the cells with the highest confidence of *Pvalb Reln* type assignment (average confidence across the ten cells on the very left: 0.88; across the other cells: 0.72).

These examples suggest that within-t-type morphoelectric variability can in some cases be related to the underlying transcriptomic variability. This is in agreement with the idea that on a fine within-family scale, both transcriptomic and morpho-electric landscapes are continuous rather than discrete.

## Discussion

Using Patch-seq, we simultaneously characterized the transcriptome, the electrophysiological properties and the morphologies of neurons from adult mouse motor cortex, providing the missing link between these modalities. We used this data set to give a description of the morphoelectric phenotypes of nearly all neural t-types in this cortical area, including some previously puzzling t-types such as *Lamp5 Lhx6* (Huang and Paul, 2019).

We found that the morpho-electric phenotype of a neuron in MOp was primarily determined by the major family of neurons it belonged to, with different families being transcriptomically as well as morpho-electrically distinct and mostly well-separated from each other (apart from some rare intermediate t-types such as *Sst Pvalb Calb2*). In contrast, within each family, variation in electrophysiological and morphological properties often appeared to be continuous across the transcriptomic landscape, without clear-cut boundaries — or gaps — between neighbouring t-types.

This seems at odds with the notion that cell types are discrete entities, a notion that is an implicit assumption behind the widespread use of clustering methods to analyze large-scale transcriptomic data sets. However, in agreement with our interpretation, several recent transcriptomic studies argued that neurons in hippocampus and striatum can be better described as forming partially continuous manifolds (Harris et al., 2018; Muñoz-Manchado et al., 2018; Stanley et al., 2019). In fact, even studies directly clustering cortical cell types have reported the prominent existence of intermediate cells with uncertain cluster assignment (Tasic et al., 2016, 2018). The goal to assemble a complete, exhaustive inventory of neural cell types might be unattainable if the types, unlike e.g. chemical elements in the periodic table, are not discrete entities. We believe that there is urgent need for theoretical work on how to conceptualize and model such a hierarchical discrete/continuous cell variability in a principled way (Zeng and Sanes, 2017).

We also found non-trivial morpho-electric variability within multiple t-types. This included variability in properties that would generally be considered to be type-defining (Zeng and Sanes, 2017). Although we cannot exclude the possibility that this variability can be attributed to some non-controlled confounding factors such as the exact spatial location of the cell within motor cortex (Cembrowski et al., 2016), there are clear cases in our data that suggest that this within-type variability is related to the within-type transcriptomic variability, in agreement with the notion of continuous phenotypic landscapes.

Developmentally, it is thought that neural diversity is generated through a combination of intrinsic genetic programs in progenitor cells, and activity-dependent and environmental factors (García et al., 2011; Dehorter et al., 2015; Wamsley and Fishell, 2017; Lim et al., 2018; Cadwell et al., 2019). It remains unclear to what extent this interplay between hard-wired genetic programs and extrinsic cues might explain our observation of discrete between family differences, and landscapes of continuous within family phenotypic variability.

Our study has several limitations. First, some t-types were covered only sparsely or even not at all. Additional experiments with more specific Cre lines could help cover some of the gaps: e.g. *L6 IT Car3* type should be accessible using Gnb4-Cre mice (Wang et al., 2019) while NP types can be targeted using another strain of the Slc17a8Cre mice (Gouwens et al., 2019). Some other t-types (e.g. *Vip Igfbp6*) might require developing new Cre lines, while some very rare putative t-types (such as *Pvalb Vipr2_1*, *Sst Pappa*, and *Sst Th_2* that together make up only 0.02% of the reference data) might not be amenable to Patch-seq study at all. Second, we found it very challenging to recover morphologies of some groups of neurons, such as e.g. deep L5 Martinotti cells with thin long axons reaching all the way to L1. This was partially due to the fact that primary motor cortex in mice is thicker than primary visual and somatosensory cortices where most studies providing anatomical descriptions of neural types have been performed (Jiang et al., 2015; Markram et al., 2015; Gouwens et al., 2019), and partially due to the RNA extraction process requiring the aspiration of the cell contents and potentially interfering with the biocytin diffusion (Cadwell et al., 2017). As a result, our study might have missed some morphological variability in this group of cells.

In parallel work, Gouwens et al. used Patch-seq to perform a similar multimodal characterization of inhibitory neurons in the visual cortex of adult mice. Our data sets are in good agreement (cf. Figure S6) and together, offer an unprecedented view at cell type variability in the neocortex. At the same time, this view is still far from complete: future studies will be needed to bring additional modalities, such as long-range projections, local connectivity, and *in vivo* functional characterization into this integrated description.

## Methods

### Animals

Experiments on adult male and female mice (*n* = 262; median age 75 days, interquartile range 64–100, full range 35–245 days, Figure S2a) were performed on wild-type (*n* = 27), Viaat-Cre/Ai9 (vesicular inhibitory amino acid transporter, encoded by the *Slc32a1* gene, *n* = 24), SstCre/Ai9 (somatostatin, *n* = 75), Vip-Cre/Ai9 (vasoactive intestinal polypeptide, *n* = 45), Pvalb-Cre/Ai9 (parvalbumin, *n* = 76), Npy-Cre/Ai9 (neuropeptide Y, *n* = 2), Vipr2-Cre/Ai9 (vasoactive intestinal peptide receptor 2, *n* = 7) and Scl17a8-Cre/Ai9 (Vglut3, vesicular glutamate transporter 3, *n* = 6) mice. Numbers above refer to mice from which the sequencing data were successfully obtained. Procedures for mouse maintenance and mouse surgeries were performed according to protocols approved by the Institutional Animal Care and Use Committee (IACUC) of Baylor College of Medicine.

The Viaat-Cre line was generously donated by Huda Zoghbi (Baylor College of Medicine). The other Cre and reporter lines were purchased from the Jackson Laboratory: Sst-Cre (stock #013044), Vip-Cre (stock #010908), Pvalb-Cre (stock #008069), Vipr2-Cre (stock #031332), Slc17a8-Cre (stock #028534), Npy-Cre (stock #027851), Ai9 reporter (stock #007909).

### Slice preparation

The MOp brain slices were obtained following previously described protocols (Jiang et al., 2015; Scala et al., 2019). Briefly, the animals were deeply anesthetized using 3% isoflurane and decapitated. The brain was rapidly removed and collected into cold (0–4 °C) oxygenated NMDG (*N* - Methyl-D-glucamine) solution containing 93 mM NMDG, 93 mM HCl, 2.5 mM KCl, 1.2 mM NaH_2_PO_4_, 30 mM NaHCO_3_, 20 mM HEPES, 25 mM glucose, 5 mM sodium ascorbate, 2 mM Thiourea, 3 mM sodium pyruvate, 10 mM MgSO_4_ and 0.5 mM CaCl_2_, pH 7.35 (all from SIGMAALDRICH). 300-μm-thick coronal slices were cut using a Leica VT1200 microtome following coordinates provided in the Allen Brain Atlas for adult mouse (http://atlas.brain-map.org). The slices were subsequently incubated at 34.0 ± 0.5 °C in oxygenated NMDG solution for 10–15 minutes before being transferred to the artificial cerebrospinal fluid solution (ACSF) containing: 125 mM NaCl, 2.5 mM KCl, 1.25 mM NaH_2_PO_4_, 25 mM NaHCO_3_, 1 mM MgCl_2_, 25 mM glucose and 2 mM CaCl_2_, pH 7.4 (all from SIGMA-ALDRICH) for about 1 hour. The slices were allowed to recover in ACSF equilibrated with CO_2_/O_2_ gas mixture (5% CO_2_, 95% O_2_), at room temperature (20–25°C) for 1 hour before experiments. During the recordings, slices were submerged in a customized chamber continuously perfused with oxygenated physiological solution.

### Patch-seq recording procedures

In order to simultaneously obtain electrophysiological, morphological and transcriptomic data from the same neurons, we applied our recently developed Patch-seq protocol (Cadwell et al., 2017), with some modifications. In particular, changes were made to the internal solution to optimize its osmolarity in order to improve staining quality. RNase-free intracellular solution was prepared as follows: we dissolved 111 mM potassium gluconate, 4 mM KCl, 10 mM HEPES, 0.2 mM EGTA in RNase-free water in a 125-ml Erlenmeyer flask. We then covered the solution with aluminum foil and autoclaved it. After it cooled down, we added 4 mM MgATP, 0.3 mM Na_3_GTP, 5 mM sodium phosphocreatine, and 13.4 mM biocytin (all from SIGMA-ALDRICH). The pH was adjusted to 7.25 with RNase-free 0.5 M KOH using a dedicated pH meter (cleaned with RNase Zap and RNase-free water before each use). RNase-free water was than added to the solution in order to obtain the desired volume. After carefully checking its osmolarity (~235–240 mOSM) the solution was stored at −20 °C and used for no longer than 3 weeks.

Before each experiment, we combined 494 μL of internal solution with 6 μL of recombinant RNase inhibitor (1 U/μL, Takara) in order to increase RNA yield. The addition of the inhibitor resulted in the increase of osmolarity to the desired value of ~315–320 mOSM without a further dilution that was described in Cadwell et al. (2017). The osmolarity of the ACSF was monitored before each experiment and adjusted to be ~18–20 mOSM lower than the internal solution by adding sucrose when needed. This osmolarity difference is important to obtain slight swelling of the cell during the recording session which improves the diffusion of biocytin in the neuronal processes. All glassware, spatulas, stir bars, counters, and anything else that may come into contact with the reagents or solution were cleaned thoroughly with RNase Zap before use.

Recording pipettes (Suttern B200-116-10) of ~3–7 MΩ resistance were filled with 0.1–0.3 μL of RNase-free intra-cellular solution. The size of the pipette tip was chosen based on the target neuron size: ~3–4 MΩ pipettes were used to record large neurons (e.g. *L5 PT* excitatory neurons) while ~6–7 MΩ pipettes were used to record small cells like L1 or *Vip* interneurons.

The PatchMaster software (HEKA Elektronik) and custom Matlab scipts were used to operate the Quadro EPC 10 amplifiers and to perform online and offline analysis of the data. We used the following quality control criteria: (1)the seal resistance value > 1 GΩ before achieving whole-cell configuration; (2) the access resistance < 30 MΩ. Each neuron was injected with 600-ms-long current pulse injections starting from 200 pA and up to 1380 pA with 20 pA increment steps (in some cases stimulation was stopped before reaching 1380 pA). There were 1.3 or 1.4 s intervals between successive current pulses, depending on the used setup. For most neurons the stimulation was then repeated multiple times from the beginning. Electrophysiological traces used for the analysis were acquired between 3–5 minutes after achieving the whole-cell configuration. All electrophysiological recordings were performed at room temperature (20–25 °C).

Typically, excitatory neurons were recorded for 5–20 minutes while interneurons were recorded for 20–50 minutes in order to allow biocytin diffusion into distal axonal segments. During the recording, the access resistance was checked every three minutes in order to maintain a stable seal that would ensure successful biocytin diffusion. The resulting cDNA yield was not correlated with the hold time (Spearman correlation −0.01).

### RNA sequencing of patched cells

At the end of the recording session, cell contents were aspirated into the glass pipette by applying a gentle negative pressure (0.7–1.5 pounds per square inch) for 1–5 minutes until the size of the cell body was visibly reduced. In most cases, the cell nucleus was visibly extracted from the cell body. During the aspiration process, the cell body structure and the access resistance were constantly monitored. Special attention was taken to ensure that the seal between the pipette and the cell membrane remained intact to avoid possible contamination from the extracellular environment. After aspiration, the contents of the pipette were immediately ejected into a 0.2 mL PCR tube containing 4 μL lysis buffer, and RNA was subsequently converted into cDNA using a Smart-seq2-based protocol (Picelli et al., 2013) as described previously (Cadwell et al., 2017). The resulting cDNA libraries were screened using an Agilent Bioanalyzer 2100. Samples containing less than ~1 ng total cDNA (in the 15 μL of the final volume) or with an average size less than 1500 bp were typically not sequenced (with some occasional exceptions).

The cDNA libraries derived from each neuron were purified and 0.2 ng of the purified cDNA was tagmented using the Illumina Nextera XT Library Preparation with one fifth of the volumes stated in the manufacturer’s recommendation. Custom 8 bp index primers were used at a final concentration of 0.1 μM. The resulting cDNA library was sequenced on an Illumina NextSeq500 instrument with a sequencing setup of 75 bp single-end reads and 8 bp index reads. Samples were sequenced in batches of ~200 cells each and the investigators were blinded to the cell type of each sample during library construction and sequencing.

The sequencing data was processed using the zUMIs pipeline with the default settings (Parekh et al., 2018). Sequencing reads were aligned to the mm10 mouse reference genome using STAR version 2.5.4b (Dobin et al., 2013) and transcript assignment performed with Gencode transcript annotations, version M23. Gene expression counts were calculated using reads mapping to exonic regions. 42 184 genes, including pseudogenes and annotated non-coding segments, were detected in at least one cell. The resulting read count data were used for all transcriptomic analyses presented in this article.

### Biocytin staining and morphological reconstructions

Morphological recovery was carried out as previously described (Jiang et al., 2015; Cadwell et al., 2017; Scala et al., 2019). Briefly, after the recordings, the slices were immersed in freshly-prepared 2.5% glutaraldehyde, 4% paraformaldehyde solution in 0.1 M phosphate-buffered saline at 4 °C for at least 48 hours. The slices were subsequently processed with the avidin-biotin-peroxidase method in order to reveal the morphology of the neurons. As described previously, we took several steps to improve the staining quality of the fine axonal branches of interneurons (Jiang et al., 2015; Cadwell et al., 2017). First, we used high biocytin concentration (0.5 g / 100 ml). Second, we incubated with avidin-biotin complex and detergents at a high concentration (Triton-X100, 5%) for at least 24 hours before DAB staining.

Recovered cells were manually reconstructed using a 100X oil-immersion lens and a camera lucida system (MicroBrightField, Vermont). We aimed to reconstruct all cells that had staining of sufficient quality (axons and dendrites for the inhibitory neurons; only dendrites for the excitatory neurons), obtaining 642 reconstructions in total. In addition, we reconstructed the dendrites of 30 neurons from the *Vip* and *Scng* families that lacked sufficient axonal staining. *Vip* neurons are traditionally classified based on the dendritic morphology, so these reconstructions can inform t-type characterizations. These additional 30 reconstructions are shown, together with the main 642 reconstructions, in the Supplementary File 1.

45 sequenced cells were mistakenly recorded using a solution with a much smaller concentration of biocytin, and their morphologies could not be recovered. We made sure that the measured electrophysiological properties of these cells were not systematically different from the other sequenced cells.

Inevitably, neuronal structures can be severed as a result of the slicing procedure. We took special care to exclude reconstructions of all neurons that showed any signs of damage, lack of contrast, or poor overall staining. Consistently with previous studies, tissue shrinkage due to the fixation and staining procedures was about 10–20% (Scala et al., 2019; Jiang et al., 2015; Markram et al., 1997). This shrinkage was not compensated for in our analysis.

### Cortical thickness normalization and layer assignment

Nissl-stained slices (*n* = 15 from 2 wild-type adult mice) were used to measure normalized layer boundaries in MOp. The Nissl staining protocol was adapted from Paul et al. (2008). Briefly, mouse brain slices were mounted on slides and allowed to dry. The sections were then demyelinated, stained with a 0.1% cresyl violet-acetate (C5042, Sigma) for 30 minutes at 60 °C and further destained. The sections were then coverslipped in Cytoseal 60 (Richard Allan Scientific). For each slice we measured its total thickness from pia to white matter and the depths of the three between-layer boundaries (L1 to L2/3, L2/3 to L5, L5 to L6), based on the cortical cytoarchitecture, using Neurolucida system with 10x/20x magnification. All measurements were normalized by the respective slice thickness, and the averages over all *n* = 15 slices were used as the normalized layer boundaries (Figure S2b).

For the Patch-seq neurons, we measured soma depth and the cortical thickness of the slice using Neurolucida system. We took their ratio as the normalized soma depth, and assigned each neuron to a layer (L1, L2/3, L5, or L6) based on the Nissl-determined layer boundaries (Figure S2b). We obtained soma depth information for 1275 neurons out of 1320 (45 neurons were mistakenly recorded using a solution with insufficient biocytin concentration, and we could only measure soma depths for 2 of those; for 2 other neurons the measurements could not be carried out because the slices were lost). For the 45 neurons with missing soma depth measurements, we used the layer targeted during the recording for all layer-based analyses and visualizations (marker shapes in Figure 1c–e and Figure 4a–c, layer-restricted analysis in Figure 5 and Figure 6).

All reconstructed morphologies were normalized by the cortical thickness of the respective slice to make it possible to display several morphologies next to each other, as in Figure 2.

### T-type assignment

The t-type assignment procedure was done in two rounds. The first round was for quality control and initial assignment to a transcriptomic ‘order’ (CGE-derived interneurons, MGE-derived interneurons, or excitatory neurons) that are perfectly separated from each other with no transcriptomically intermediate cells (Tasic et al., 2018). The second round was done for assigning the cells to specific t-types.

In the first round, we mapped each Patch-seq cell to a large annotated Smart-seq2 reference data set from adult mouse cortex (Tasic et al., 2018), using the same procedure as in Scala et al. (2019). Specifically, using the exon count matrix of Tasic et al. (2018), we selected 3000 most variable genes (see below). We then log-transformed all counts with log_2_(*x* + 1) transformation and averaged the log-transformed counts across all cells in each of the 133 t-types, to obtain reference transcriptomic profiles of each t-type (133 × 3000 matrix). Out of these 3000 genes, 2664 were present in the genome annotation that we used. We applied the same log_2_(*x* + 1) transformation to the read counts of our cells, and for each cell computed Pearson correlation across the 2686 genes with all 133 t-types. Each cell was assigned to the t-type to which it had the highest correlation (Figure S4a).

Cells meeting any of the exclusion criteria described in the following were declared low quality and did not get a t-type assignment (Figure S2e): cells with the highest correlation below 0.4 (76 cells); cells that would be assigned to non-neural t-types, presumably due to RNA contamination (Tripathy et al., 2018) (12 cells); cells that would be assigned to the *Meis2* t-type that was not adequately represented in the datasets from Yao et al. (in preparation) (2 cells); cells with the highest correlation less than 0.02 above the maximal correlation in other transcriptomic orders (2 cells). The remaining 1228 cells passed the quality control and entered the second round.

In the second round, cells were independently mapped to the seven transcriptomic datasets from Yao et al. (in preparation) obtained from mouse MOp (Figure S4b). The mapping was done only to the t-types from the transcriptomic order identified in the first round, using 500 most variable genes in that dataset for that transcriptomic order (so using 21 sets of 500 most variable genes). Gene selection was performed as described below, and t-type assignment was done exactly as described above. Across the 21 reference subsets, 472–494 most variable genes were present in the genomic annotation used here, and were used for the t-type assignment.

We used bootstrapping over genes to assess the confidence of each t-type assignment. For each cell and for each reference dataset, we repeatedly selected a bootstrap sample of genes (i.e. the same number of genes, selected randomly with repetitions) and repeated the mapping. This was done 100 times and the fraction of times the cell mapped to each t-type was taken as the t-type assignment confidence for that t-type (Figure S4c). The confidences obtained with seven reference data sets agreed well with each other (Figure S2i) and were averaged to obtain the consensus confidence. Finally, the cell was assigned to the t-type with the highest consensus confidence.

Six cells were assigned to an excitatory t-type, despite having clearly inhibitory firing, morphology, and/or soma depth location (such as L1). The most likely cause was RNA contamination from excitatory cells that are much more abundant in the mouse cortex. These six cells were excluded from all analyses and visualizations (as if they did not pass the transcriptomic quality control). In addition, one cell was likely located outside of MOp, based on the slice anatomy, and was excluded as well. The final number of cells with t-type assignment was 1221.

### Selection of most variable genes

Several steps of our analysis required selecting a set of the most variable genes in a given transcriptomic data set. We always selected a fixed predefined number of genes (such as 500, 1000, or 3000).

To select the most variable genes, we found genes that had, at the same time, high non-zero expression and high probability of near-zero expression (Andrews and Hemberg, 2019). Our procedure is described in more detail elsewhere (Kobak and Berens, 2019). Specifically, we excluded all genes that had counts of at least *c*_min_ (for Patchseq and Smart-seq2: *c*_min_ = 32; for 10x: *c*_min_= 1) in fewer than 10 cells. For each remaining gene we computed the mean log_2_ count across all counts that were larger than *c*_min_ (non-zero expression, *μ*) and the fraction of counts that were smaller than *c*_min_ (probability of near-zero expression, *τ*). Across genes, there was a clear inverse relationship between *μ* and *τ*, that roughly followed exponential law

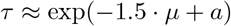

for some horizontal offset *a*. Using a binary search, we found a value *b* of this offset that yielded the desired number of genes with

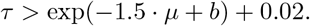

### T-SNE visualization of the transcriptomic data

T-SNE embeddings (Maaten and Hinton, 2008) of the three subsets of the single-cell 10x v2 data set from Yao et al. (in preparation) (Figure 1c–e) were constructed using the same 500 most variable genes that were also used for t-type assignment (see above). The counts were converted to counts per million (CPM), log_2_(*x* +1)-transformed, and reduced to 50 principal components. The resulting *n* × 50 matrix was used as input to t-SNE. We used FIt-SNE 1.1.0 (Linderman et al., 2019) with learning rate *n/*12 and scaled PCA initialisation (Kobak and Berens, 2019). Perplexity was left at the default value of 30 for both inhibitory subsets and increased to 100 for the excitatory subset. All other parameters were left at default values.

The embedding of the full data set (Figure S5) was constructed using the same 3000 most variable genes as were used for t-type assignment (as above). Counts were converted to CPMs, log_2_(*x* + 1)-transformed, and reduced to 50 PCs. For t-SNE, we used downsampling-based initialisation (Kobak and Berens, 2019). Briefly, the data set was randomly downsampled to 25 000 cells and embedded using the learning rate *n/*12, perplexity combination of 30 and *n/*100, and scaled PCA initialisation. The remaining cells were positioned in the median embedding location of their 10 nearest neighbours (based on Euclidean distance in the high-dimensional space). The resulting embedding, scaled to have standard deviation 0.0001, was used as initialization for t-SNE with default perplexity 30 and learning rate *n/*12.

To position Patch-seq cells on a reference t-SNE embedding, we used the procedure from Kobak and Berens (2019). Briefly, each cell was positioned at the median embedding location of its 10 nearest neighbours, based on Pearson correlation distance in the high-dimensional space. All counts were log_2_(*x* + 1)-transformed and correlations were computed across the same genes that were used for t-type assignments (see above).

### Extraction of electrophysiological features

28 electrophysiological properties of the neurons were automatically extracted based on the raw membrane voltage traces (Figure S12) using Python scripts from the Allen Software Development Kit (https://github.com/AllenInstitute/AllenSDK) with some modifications to account for our experimental paradigm (https://github.com/berenslab/EphysExtraction).

For each hyperpolarizing current injection, the *resting membrane potential* was computed as the mean membrane voltage during 100 ms before stimulation onset and the *input resistance* as the difference between the steady state voltage and the resting membrane potential, divided by the injected current value (we took the average voltage of the last 100 ms before stimulus offset as steady state). The median of these values over all hyperpolarizing traces was taken as the final resting membrane potential and input resistance respectively.

To estimate the *rheobase* (the minimum current needed to elicit any spikes), we used robust regression (random sample consensus algorithm, as implemented in `sklearn.linear_model.RANSACRegressor`) of the spiking frequency onto the injected current using the five lowest depolarizing currents with non-zero spike count (if there were fewer than five, we used those available). The point where the regression line crossed the *x*-axis gave the rheobase estimate (Figure S12). We restricted it to be between the highest injected current eliciting no spikes and the lowest injected current eliciting at least one spike. In case the regression line crossed the *x*-axis outside of this interval, the first current step eliciting at least one spike was used.

The *action potential (AP) threshold, AP amplitude, AP width, afterhyperpolarization (AHP), afterdepolarization (ADP)*, the first AP *latency*, and the *upstroke-to-downstroke ratio (UDR)* were computed as illustrated in Figure S12, using the very first AP fired by the neuron. AP width was computed at the AP half-height. UDR refers to the ratio of the maximal membrane voltage derivative during the AP upstroke to the maximal absolute value of the membrane voltage derivative during the AP downstroke.

The *interspike interval (ISI) adaptation index* for each trace was defined as the ratio of the second ISI to the first one. The *ISI average adaptation index* was defined as the mean of ISI ratios corresponding to all consecutive pairs of ISIs in that trace. For both quantities we took the median over the five lowest depolarizing currents that elicited at least three spikes (if fewer than five were available, we used all of them). *AP amplitude adaptation index* and *AP amplitude average adaptation index* were defined analogously to the two ISI adaptation indices, but using the ratios of consecutive AP amplitudes (and using the median over the five lowest depolarizing currents that elicited at least two spikes).

The *maximum number of APs* refers to the number of APs emitted during the 600 ms stimulation window of the highest firing trace. The *spike frequency adaptation (SFA)* denotes the ratio of the number of APs in the second half of the stimulation window to the number of APs in the first half of the stimulation window of the highest firing trace. If the highest firing trace had fewer than five APs, SFA was not defined. Here and below the highest firing trace corresponds to the first depolarising current step that showed the maximum amount of APs during the current stimulation window (after excluding all stimulation currents for which at least one AP was observed in 100 ms before or in 200 ms after the stimulation window; see below).

The *membrane time constant* (tau) was computed as the time constant of the exponential fit to the membrane voltage from the stimulation onset to the first local minimum (we took the median over all hyperpolarizing traces). Three further features described the sag of the first (the lowest) hyperpolarization trace. The *sag ratio* was defined as the sag trough voltage (average voltage in a 5 ms window around the sag trough) difference from the resting membrane potential, divided by the steady state membrane voltage difference from the resting membrane potential. The *sag time* was defined as the time period between the first and the second moments the membrane voltage crosses the steady state value after the stimulation onset. The *sag area* refers to the absolute value of the integral of the membrane voltage minus the steady-state voltage during the sag time period (Figure S12). If the sag trough voltage and the steady state voltage differed by less than 4 mVs, the sag time and sag area were set to zero.

The *rebound* was defined as the voltage difference between the resting membrane potential and the average voltage over 150 ms (or whatever time remained until 300 ms after the stimulation offset) after rebound onset, which we identified as the time point after stimulation offset when the membrane voltage reached the value of the resting membrane potential. If the membrane voltage never reached the resting membrane potential during the 300 ms after the stimulation offset, the rebound was set to zero. The *rebound number of APs* was defined as the number of APs emitted during the same period of time. Both rebound features were computed using the lowest hyperpolarization trace.

The *ISI coefficient of variation (CV)* refers to the standard deviation divided by the mean of all ISIs in the highest firing trace. Note that a Poisson firing neuron would have ISI CV equal to one. The *ISI Fano factor* refers to the variance divided by the mean of all ISIs in the highest firing rate. The *AP CV* and *AP Fano factor* refer to the CV and the Fano factor of the AP amplitudes in the highest firing trace.

The *burstiness* was defined as the difference between the inverse of the smallest ISI within a detected burst and the inverse of the smallest ISI outside of bursts, divided by their sum. We took the median over the first five depolarizing traces. We relied on the Allen SDK code to detect the bursts. Briefly, within that code a burst onset was identified whenever a ‘detour’ ISI was followed by a ‘direct’ ISI. Detour ISIs are ISIs with a non-zero ADP or a drop of at least 0.5 mVs of the membrane voltage after the first AP terminates and before the next one is elicited. Direct ISIs are ISIs with no ADP and no such drop of membrane voltage before the second AP. A burst offset was identified whenever a direct ISI was followed by a detour ISI. Additionally, bursts were required to contain no ‘pause-like’ ISIs, defined as unusually long ISIs for that trace (see Allen SDK for the implementation details).

Some neurons (in particular neurogliaform cells) started emitting APs before and after the current stimulation window, after the stimulation currents exceeded certain amount. To quantify this effect, we defined *wildness* as the difference in the number of APs between the highest firing trace (possibly showing APs before or after stimulation window) and the highest firing trace as defined above (without any APs outside the stimulation window). For most neurons, wildness was equal to zero.

For all statistical analysis we used 16 features out of the extracted 28, excluding features that were equal to zero for many cells (afterdepolarization, burstiness, rebound number of APs, sag area, sag time, wildness), two Fano factor features that were highly correlated with the corresponding coefficient of variation features (AP Fano factor, ISI Fano factor), features that had very skewed distributions (AP amplitude average adaptation index, ISI average adaptation index), features that could not be extracted for some of the cells (spike frequency adaptation), and features that we considered potentially unreliable (latency). Three features were log-transformed to make their distribution more Gaussian-like: AP coefficient of variation, ISI coefficient of variation, ISI adaptation index.

### Extraction of morphological features

Reconstructed morphologies were converted into the SWC format using `NLMorphologyConverter 0.9.0` (http://neuronland.org) and further analyzed in Python. Each cell was soma-centered in the *x* (slice width) and *y* (slice depth) dimensions, and aligned to pia in the *z* (cortical depth) dimension so that *z* = 0 corresponded to pia. All neurites were smoothed in the slice depth dimension (*y*) using a Savitzky-Golay filter of order 3 and window length 21, after resampling points to have maximally 1 *μ*m spacing. For further analysis we computed two different feature representations of each cell: the normalized *z*-profile and a set of morphometric statistics (Scala et al., 2019; Gouwens et al., 2019; Laturnus et al., 2019).

To compute the normalized *z*-profile, we divided all the coordinates of the neuronal point cloud by the thickness of the respective cortical slice, so that *z* = 1 corresponded to the white matter border. We projected this point cloud onto the *z*-axis and binned it into 20 equal-sized bins spanning [0, 1]. The resulting histogram describes a neuron’s normalised depth profile perpendicular to the pia. For the purposes of downstream analysis, we treated this as a set of 20 features. The *z*-profiles were separately computed for axons and dendrites.

Morphometric statistics were separately computed for the dendritic and the axonal neurites to quantify their arborization shape and branching patterns. For the excitatory neurons, several additional morphometric statistics were computed for the apical dendrites, where apical dendrite was operationally defined as the dendrite with the longest total path length. We further used two ‘somatic’ features, normalized soma depth and soma radius. We did not use any features measuring morphological properties in the slice depth (*y*) direction due to possible slice shrinkage artefacts. We did not use any axonal features for the excitatory cells because only a small part of the axon could typically be reconstructed. For the inhibitory cells, where dendrite and axon can both be fully recovered, we included some measures of the dendritic and axonal overlap. The full list of morphometric statistics is given in Table S1.

We extracted a set of 75 features, of which 40 were defined for excitatory neurons and 58 for inhibitory neurons, and processed the data for excitatory and inhibitory neurons separately. In each case, we excluded features with coefficient of variation below 0.25 (among the features with only positive values). This procedure excluded 5 features for the excitatory and 10 features for the inhibitory cells. The distributions of the remaining features were visually checked for outliers and for meaningful variation between transcriptomic types, leading to a further exclusion of 3 and 4 features for the excitatory and the inhibitory cells respectively. The full list of excluded features is given in Table S1. The resulting set of morphometric statistics used for further analysis consisted of 32 features defined for the excitatory neurons and 48 features defined for the inhibitory neurons.

### Reduced-rank regression

For the reduced-rank regression (RRR) analysis (Kobak et al., 2019) we used 16 electrophyiological features and all 1213 cells for which values for all 16 features and a t-type assignment could be computed. Electrophyiological features were standardized. Transcriptomic counts were converted to CPM, log_2_(*x*+1)-transformed, and then standardized. We selected the 1000 most variable genes and only used those for the RRR analysis.

Briefly, RRR finds a linear mapping of gene expression levels to a low-dimensional latent representation, from which the electrophysiological features are then predicted with another linear transformation (for mathematical details, see Kobak et al. (2019)). The model employs sparsity constraints in form of elastic net penalty to select only a small number of genes. For Figure 3 we used a model with rank *r* = 5, zero ridge penalty (*α* = 1), and lasso penalty tuned to yield a selection of 25 genes (*λ* = 0.45). Crossvalidation (Figure S8) was done using 10 folds, elastic net *α* values 0.5, 0.75, and 1.0, and *λ* values from 0.2 to 6.0.

The plots shown in Figure 3a and Figure 3b are called bibiplots because they combine two biplots: the left biplot shows a mapping of gene expression levels on the two latent dimensions; the right biplot shows the same mapping of electrophysiological features. To illustrate the meaning of the latent dimensions, each biplot combines the resulting scatter plots with lines showing how original features are related to the latent dimensions. Specifically, we computed the correlations of individual genes or electrophysiological properties with the latent dimensions and visualized these correlations as lines on the biplot. The circle shows the maximal possible correlation; only lines longer than 0.4 times the circle radius were shown in Figure 3.

For the model based on ion channel genes, we obtained the list of 328 ion channel genes from https://www.genenames.org/data/genegroup/#!/group/177 and used 293 of them that had non-zero expression in at least 10 of our cells. We used rank *r* = 5, *α* = 1, and *λ* tuned to yield 25 genes (*λ* = 0.325), as above.

### T-SNE visualization of the morpho-electric phenotypes

For the t-SNE visualization (Maaten and Hinton, 2008) of the electrophysiological phenotypes, we used 16 features as described above and all *n* = 1311 cells that had values for all 16 features. All features were standardized across this set of cells and transformed with PCA into a set of 16 PCs. We scaled the PCs by the standard deviation of PC1. We used the t-SNE implementation from `scikit-learn` Python library with the default perplexity (30), early exaggeration 4 (the default value 12 can be too large for small datasets), and scaled PCA initialisation (Kobak and Berens, 2019). Figure 4a only shows *n* = 1218 cells that had a t-type assignment.

For the t-SNE visualization of the morphological phenotypes, we combined morphometric statistics with the normalized *z*-profiles. The pre-processing, including PCA, was done separately for the excitatory and for the inhibitory neurons because they used different sets of morphometric statistics (see above). Only neurons with assigned t-type were used for this analysis. Two inhibitory neurons were left out because some of the morphometric statistics could not be extracted due to insufficient dendritic recovery; this left 367 inhibitory neurons (with 48 morphometric features) and 268 excitatory neurons (with 32 morphometric features). All features were standardized and each set was reduced to 20 PCs. We scaled the PCs by the standard deviation of the respective PC1, to make the inhibitory and the excitatory PCs have comparable variances.

We used dendritic *z*-profiles for the excitatory neurons and axonal *z*-profiles for the inhibitory neurons. We reduced each set to 5 PCs, discarded PC1 (it was strongly correlated with the normalized soma depth and made the resulting embedding strongly influenced by the soma depth), and scaled the PCs by the standard deviation of the respective PC2. We stacked the 20 scaled morphometric PCs and the 4 scaled *z*-profile PCs to get a combined 24-dimensional representation, separate for the excitatory and for the inhibitory neurons. We then combined these representations into one block-diagonal 48-dimensional matrix. This procedure makes the excitatory and the inhibitory populations both have zero mean. To prevent this overlap, we added a small constant value of 0.3 to the excitatory block-diagonal block, leading to the strong excitatory/inhibitory separation in Figure 4b. The t-SNE was performed exactly as described above.

For the t-SNE visualization of the morpho-electrical landscape, we stacked together the 48-dimensional morphological representation and the 16-dimensional electro-physiological representation obtained above, using only cells that had all morphological and all electrophysiologcal features (*n* = 627). We multiplied the electro-physiological block by 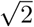 to put its total variance on a similar scale (it only consisted of one set of scaled PCs, whereas the morphological representation consisted of two sets of scaled PCs: morphometrics and *z*-profiles). The resulting 64-dimensional morpho-electrical representation was used for t-SNE, exactly as described above.

### kNN classification of transcriptomic families

To classify neurons into transcriptomic families based on the electrophysiological, morphological, or combined features (Figures 4d, 6a), we used a kNN classifier with *k* = 10 and Euclidean distance metric (taking the majority family among the *k* nearest neighbours). This is effectively a leave-one-out cross-validation procedure. For each data modality we took the exact same data representation that was used for computing t-SNE embeddings (Figure 4a–c), see above. Note that t-SNE algorithm is also based on nearest neighbors and makes all close neighbors attract each other in the embedding. We chose the kNN classifier as a simple but versatile non-parametric classifier that is directly related to the t-SNE embeddings. We did not use the *Sncg* and NP families due to insufficient coverage in our data set (Figure 1).

Figure 4d shows the fraction of cells from each family that got classified into each family. Figure 6a shows fractions of cells from each t-type that got classified into each family. For morphological and combined features, it shows fractions of cells from the majority layer of each t-type. E.g. *Pvalb Reln* type occurred most often in L5, so only cells from that layer were taken for that type. Only t-types with at least 10 cells (or at least 10 layer-restricted cells) are shown.

### Within-family analysis

To study the relationship between transcriptomic and electrophyiological distances between pairs of t-types (insets in Figure 5a,b, Figure S10), we took all t-types with at least 10 cells. For each pair of t-types, transcriptomic distance was computed as the Pearson correlation between the average log_2_(*x*+1)-transformed gene counts in the single-cell 10x v2 data from Yao et al. (in preparation). 1000 most variable genes across all neural types were used for the Figure S10 and 500 most variable genes across the respective family were used for the Figure 5a,b. Electrophysiological distance was computed as the Euclidean distance between the average feature vectors.

Figure 5c used the soma depth distance, computed as the absolute value of the difference between the average normalized soma depths. Here we used all t-types with at least 5 cells.

### T-type variability analysis

The normalized total variance in Figure 6b was computed as follows. For each modality, we took the exact same data representation that was used for computing t-SNE embeddings (Figure 4a–c), see above. For each t-type (or layer-restricted t-type, see above), we took the sum of its variances in all dimensions as the total variance and divided by the sum of variances in all dimensions across the whole data set:

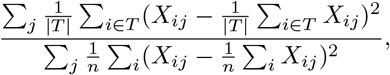

where *X*_ij_ is a value of feature *j* of cell *i*, *n* is the total number of cells, and *T* is the set of cell numbers belonging to the given t-type. The value 0 indicates that all cells from this t-type have exactly identical features. The value 1 indicates that there is as much variance in this one t-type as in the whole data set. Only t-types with at least 10 cells (or at least 10 layer-restricted cells) are shown in Figure 6b.

To provide a sensible baseline for the range of possible normalized total variances in a population of morpho-electrically homogeneous types, we used a clustering analysis. For the cells of all the *K* t-types (or layer-restricted t-types) with at least 10 cells in a given panel, we used the k-means algorithm to cluster them into *K* clusters, reasoning that these clusters should be as homogeneous as possible given the variability in our data set. We used the k-means implementation from `scikit-learn` with default parameters. We then computed the normalized total variance of each cluster as described above. Grey shading in Figure 6b shows the interval between the minimum and the maximum cluster variances. Note that the k-means algorithm is directly minimizing within-cluster total variances.

We used the entropies of a Leiden clustering (Traag et al., 2019) as an alternative way to approach the same question. For each modality, using the exact same data representation as above, we constructed its kNN graph with *k* = 10 and clustered it using the Leiden algorithm as implemented in the Python package `leidenalg` with `RBConfigurationVertexPartition` quality function and resolution parameter manually tuned to produce roughly the same number of clusters for each modality as in Gouwens et al. (2019) (Figure S11). For each t-type (or layer-restricted t-type) we then measured the entropy of the distribution of electrophysiological/morphological cluster ids, after randomly subsampling the t-type to 10 cells. Subsampling was done to eliminate a possible bias due to the t-type abundance. The whole procedure was repeated 100 times with different random seeds for the Leiden clustering and for the subsampling.

## Acknowledgements

We thank Ben Dichter for his help with converting electrophysiologial recordings into the NWB format.

This work was supported by the National Institute of Mental Health of the National Institutes of Health under Award Number U19MH114830, the Deutsche Forschungsgemeinschaft through a Heisenberg Professorship (BE5601/4-1), the Cluster of Excellence “Machine Learning — New Perspectives for Science” (EXC 2064, project number 390727645) and the Collaborative Research Center 1233 “Robust Vision” (project number 276693517), the German Federal Ministry of Education and Research (FKZ 01GQ1601 and 01IS18039A). The content is solely the responsibility of the authors and does not necessarily represent the official views of the National Institutes of Health.

## Author contributions

FS, DK, RS, PB and AST designed the study; FS and MB performed Patch-seq experiments; SAM performed Nissl stainings; FS and EM performed neuronal reconstructions aided by JRC; JRC prepared cDNA libraries and performed DAB stainings; LH performed sequencing and initial bioinformatic analysis under the supervision of RS; ZY and HZ shared the reference data sets; CRC and XJ helped adjusting the Patch-seq procedure; YB analyzed electrophysiological data; SL analyzed morphological data; DK analyzed transcriptomic data; DK and PB supervised data analysis; JRC and ZT sustained animal colonies and provided experimental support; FS, DK, YB, LH, SL, RS, PB and AST discussed the results; DK prepared the figures; DK and PB wrote the manuscript with input from FS, YB, SL, LH, CRC, RS, and AST.

## Competing interests

The authors declare no competing interests.

## Data availability

All data will be made available using dedicated BICCN online archives in February–April 2020.

## Code availability

The analysis code will be made available at https://github.com/berenslab/mini-atlas in February 2020.

## Supplementary Figures

**Figure S1:**
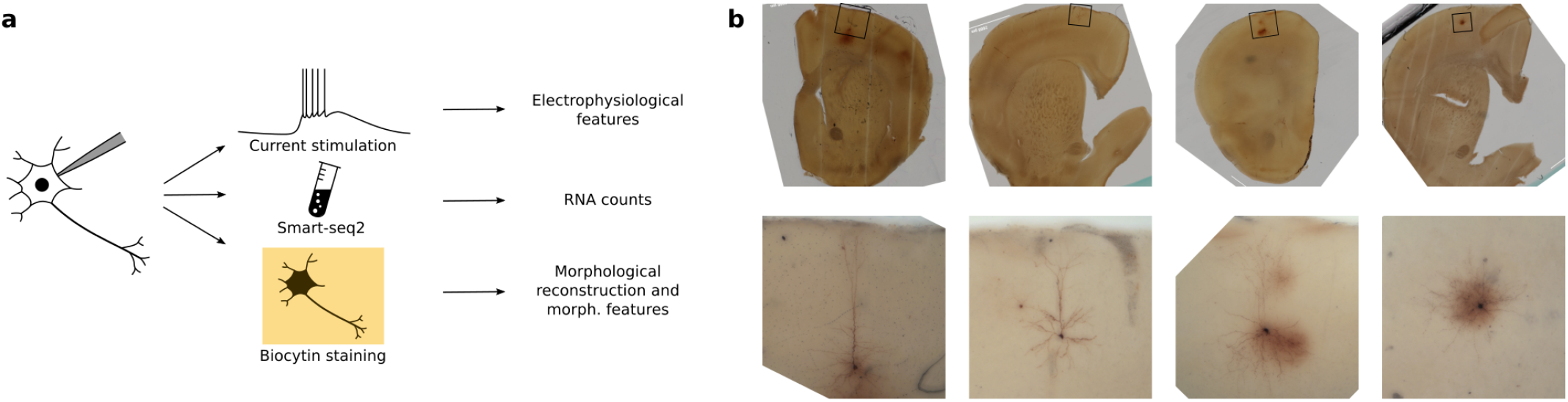
Patch-seq protocol. **(a)** Patch-seq combines electrophyiological recordings, RNA sequencing using Smart-seq2, and biocytin staining in the same cell. **(b)** Four exemplary slice images. Top: an image of the whole slice using 4x magnification. Bottom: a flattened 3D image stack using 20x magnification. From left to right: L5 PT neuron, L2/3 IT neuron, L5 *Sst* neuron, L5 *Pvalb* neuron.

**Figure S2:**
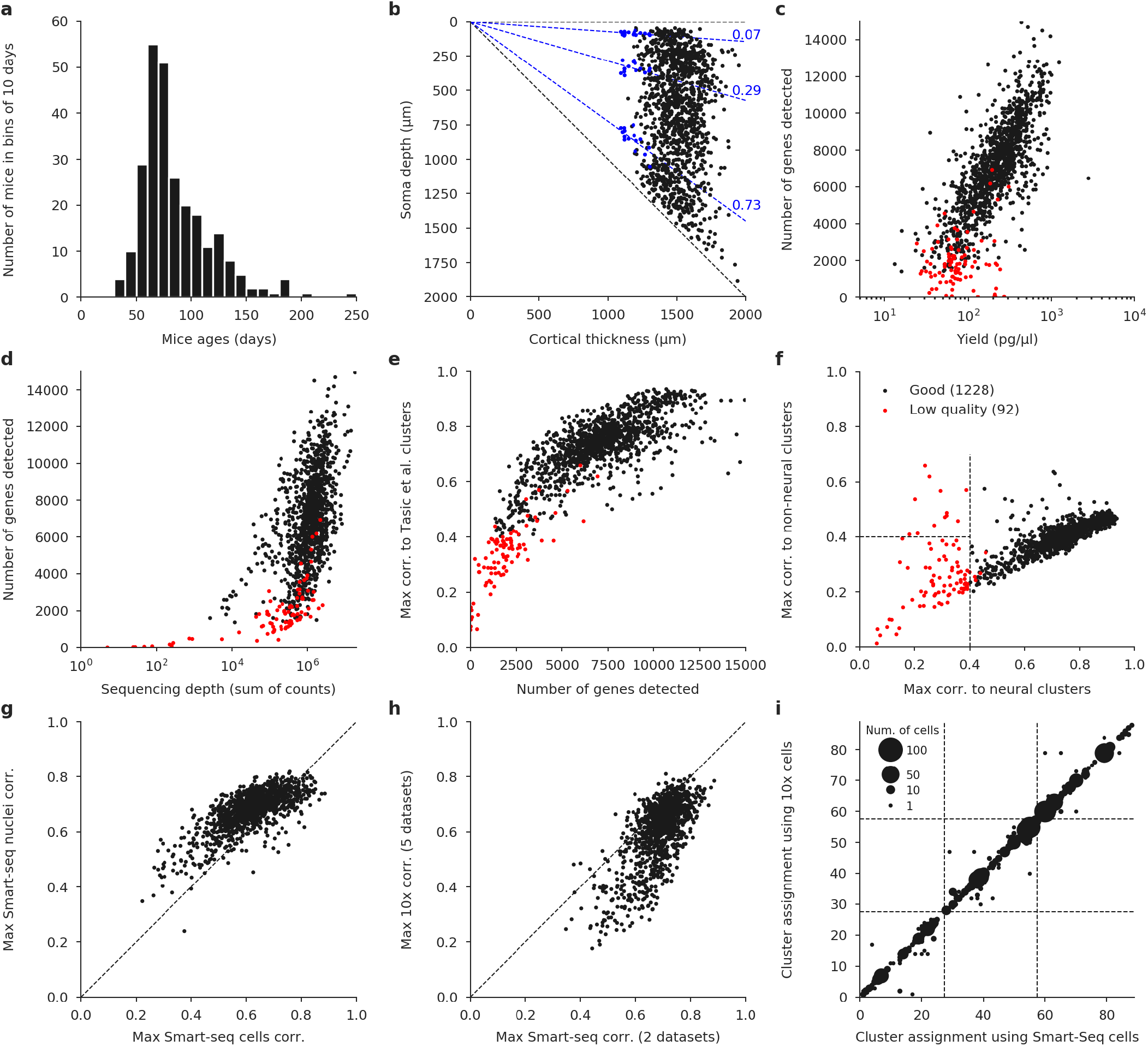
Quality control. **(a)** Age distribution of the mice used in the experiments. Median: 75 days. **(b**) Soma depths of all cells and cortical thickness of the corresponding slices. Dashed lines show layer boundaries, based on the Nissl-stained slices (measured layer boundaries shown as blue points). All soma depths were normalized by dividing them by the cortical thickness. **(c)** Relationship between cDNA yield and the number of genes detected after sequencing. Cells with very low yield were typically not sequenced. Red: cells eventually failing quality control. **(d)** Relationship between sequencing depth (total number of reads) and the number of detected genes (number of genes with non-zero counts). **(e)** Relationship between the number of detected genes and the maximal correlation to clusters from Tasic et al. (2018). Cells with maximal correlation below 0.4 were declared low quality. **(f)** Relationship between the maximal correlation across neural clusters and the maximal correlation across non-neural clusters from Tasic et al. (2018). Cells with maximal neural correlation below 0.4 were declared low quality. See Methods for additional QC creiteria. **(g)** Maximal correlations using single-cell and single-nuclei Smart-seq2 reference data sets (Yao et al., in preparation). **(h)** Maximal correlations using Smart-seq2 reference data sets (maximum across cell types and across two data sets) and using 10x reference data sets (maximum across cell types and across five data sets). **(i)** T-type assignment using single-cell Smart-seq2 reference data set and using single-cell 10x v2 reference data set. All points are on the integer grid; marker size shows the number of cells at the corresponding location. Dashed lines separate three taxonomic orders: CGE-derived interneurons, MGE-derived interneurons, and excitatory cells. The mapping was done within each order, so there cannot be any cells outside of the diagonal blocks.

**Figure S3:**
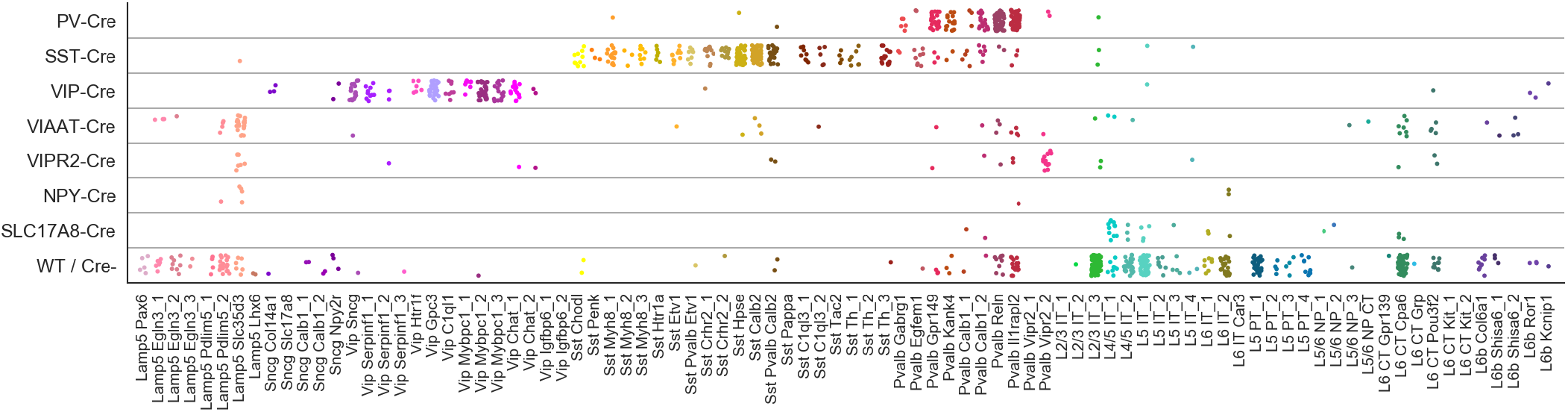
T-types assigned to cells collected in mice from different Cre lines. ‘WT / Cre-’ stands for cells from any Cre line that were not labeled with a fluorescent indicator, or for the cells patched in wild type mice. 1221 cells shown.

**Figure S4:**
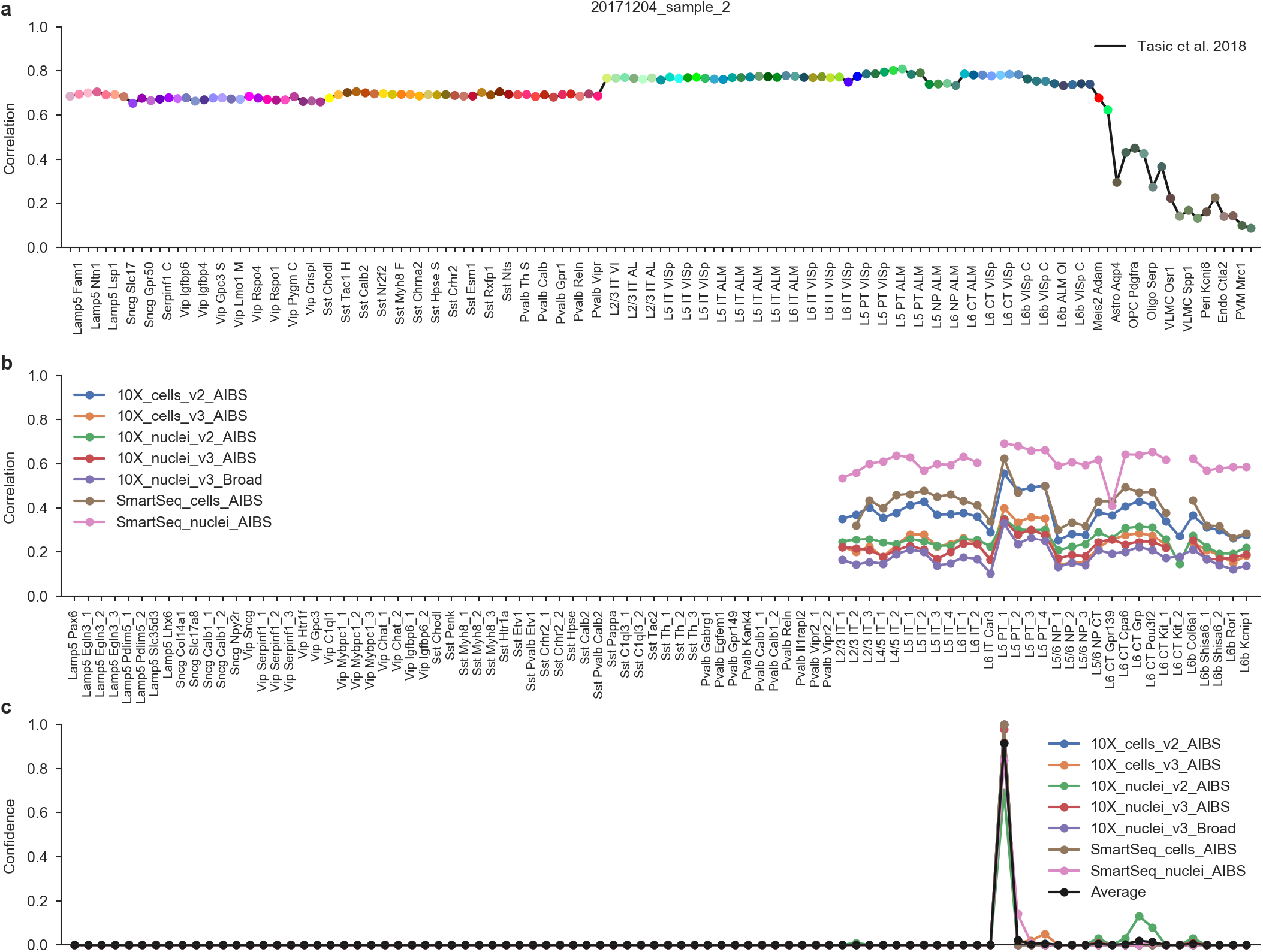
T-type assignment procedure. All panels show one particular Patch-seq cell. **(a)** Correlations to the mean log expression of all t-types from Tasic et al. (2018), using 3000 most variable genes. Maximum correlation is to the excitatory neurons. **(b)** Correlations to all excitatory t-types from Yao et al. (in preparation) using all seven reference data sets and 500 most variable genes. **(c)** T-type assignment confidences for all seven datasets, obtained via bootstrapping over genes. The average confidence is shown in black. The mode of the average confidence was taken as the final t-type.

**Figure S5:**
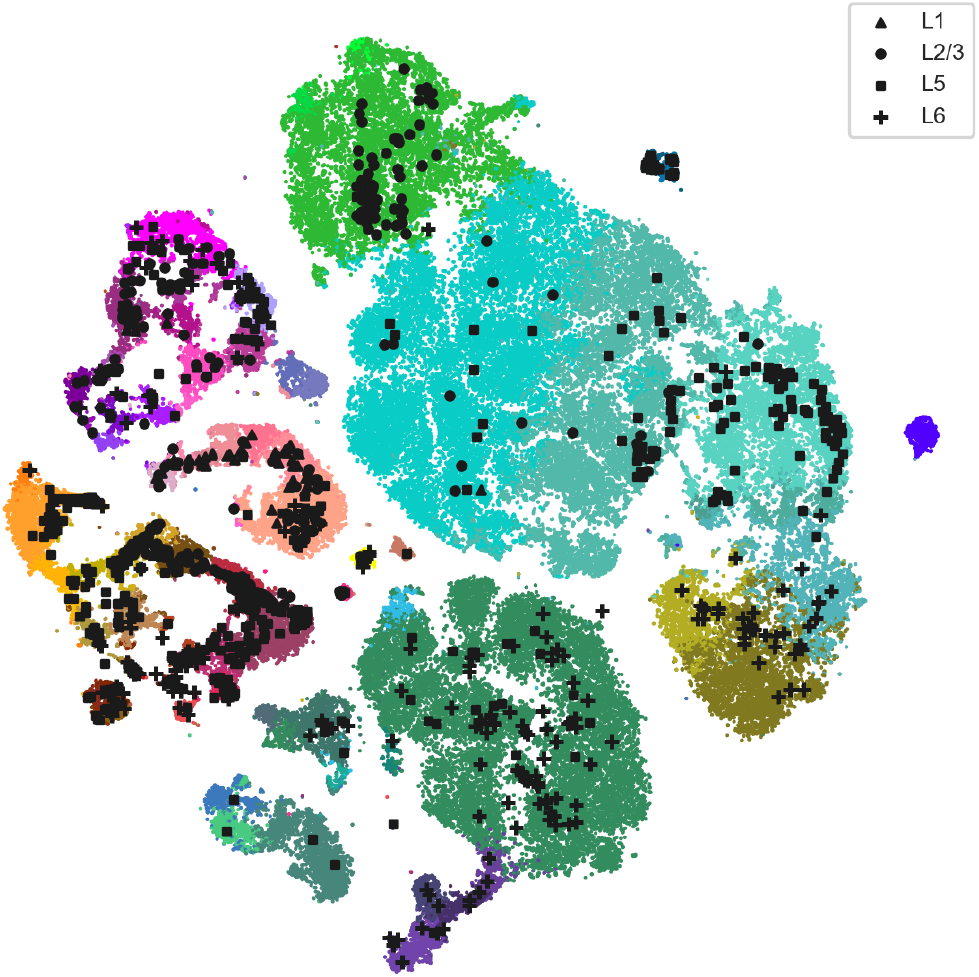
Full t-SNE of MOp neurons. Combined t-SNE of all neurons from the single-cell 10x v2 reference data set (Yao et al., in preparation), with our cells overlayed as in Figure 1c–e. Sample size 121 423, perplexity 30, downsampling-based initialization (Kobak and Berens, 2019).

**Figure S6:**
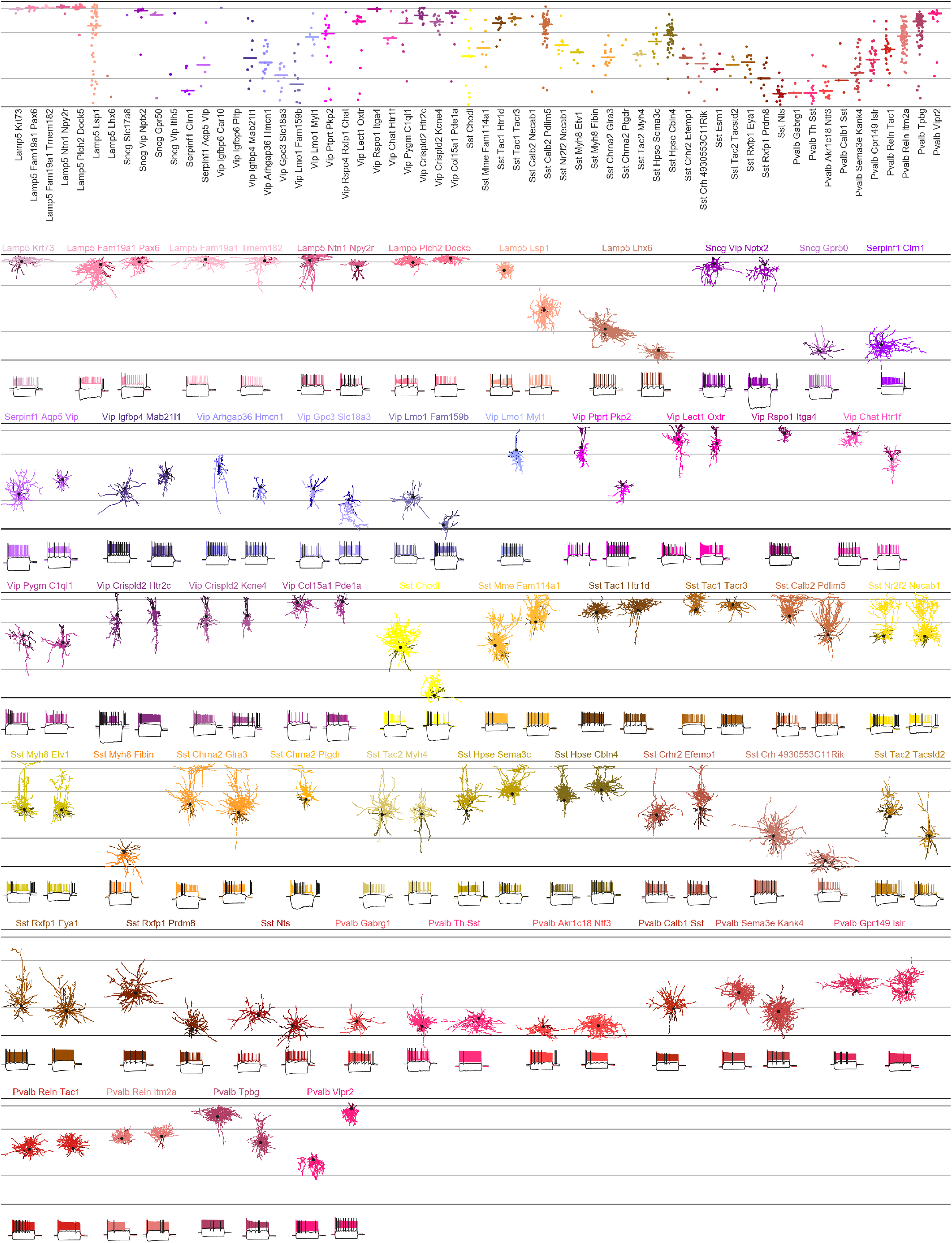
Interneurons assigned to the Tasic et al. 2018 t-types. This is an exact analogue of Figures 1b and 2 using inhibitory t-types from Tasic et al. (2018). It allows the direct comparison with the results from the parallel work by Gouwens et al. (in preparation). We used the same neurons as in Figure 2 whenever possible. 97 neurons in 53 t-types.

**Figure S7:**
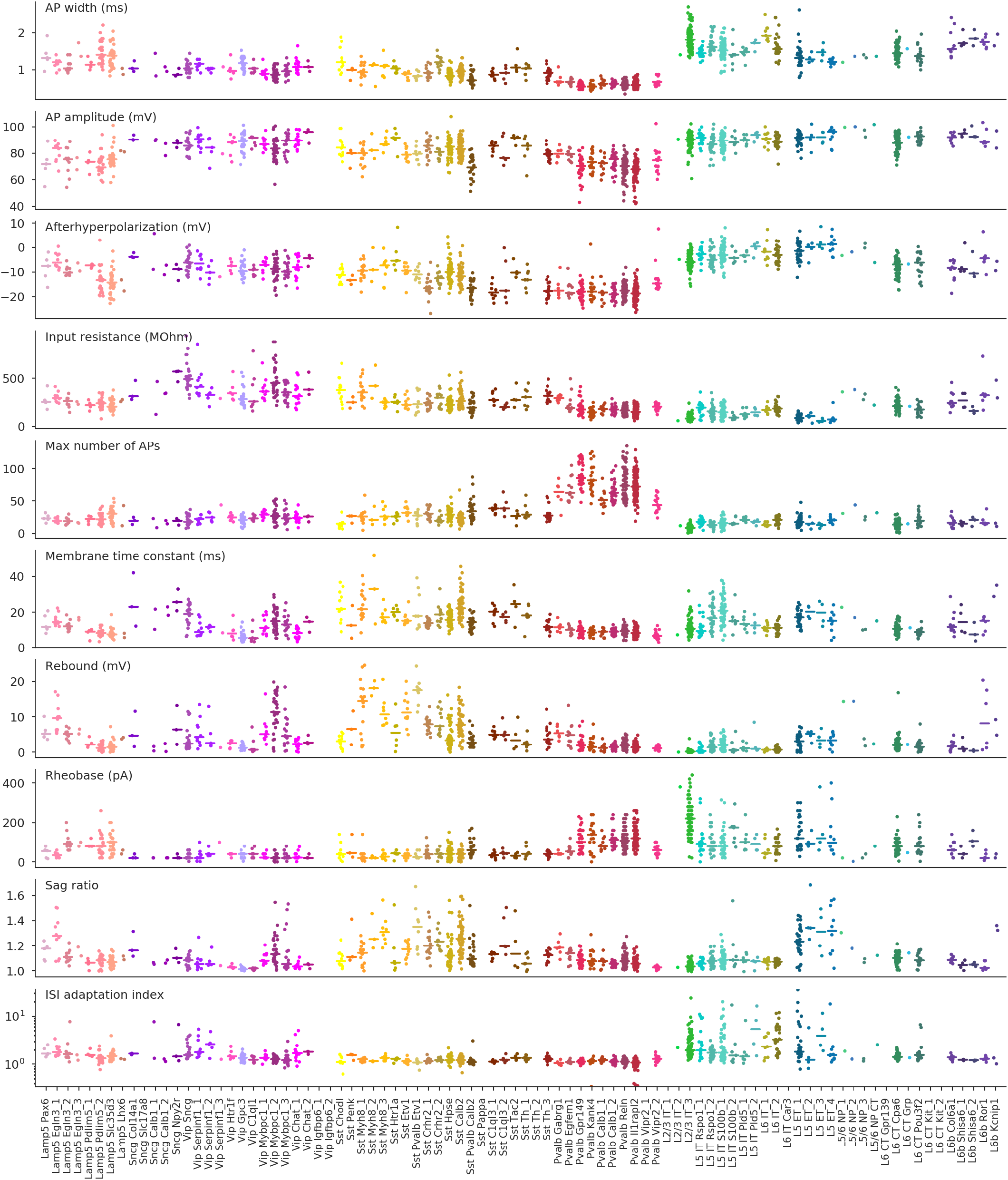
Distribution of electrophysiological features. The ten most important electrophyiological features are shown for all cells across all t-types. For t-types with at least three cells, horizontal lines show median value.

**Figure S8:**
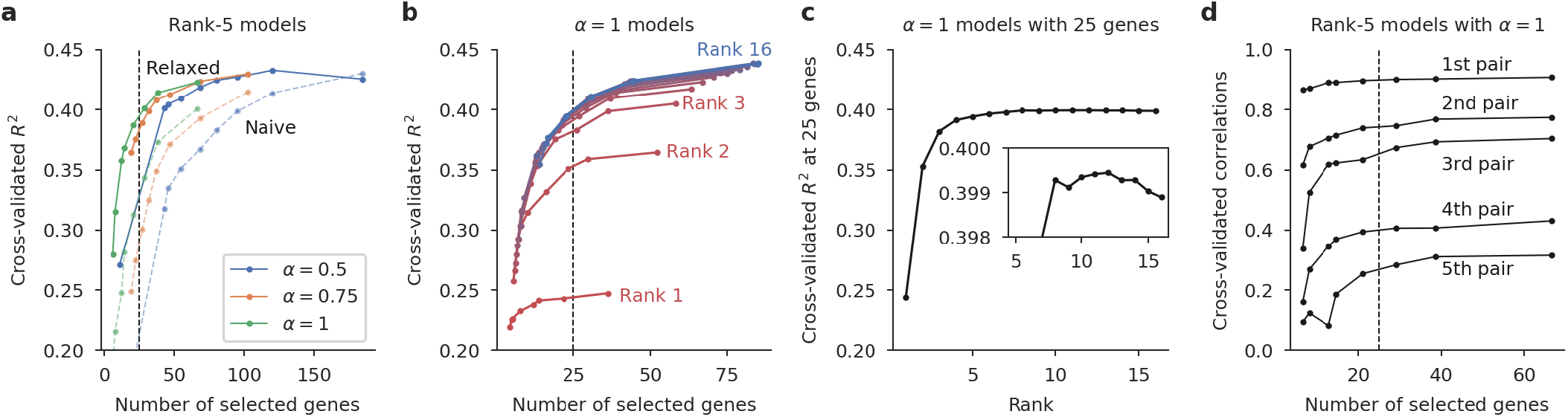
Cross-validation for the reduced-rank regression models. **(a)** Cross-validated *R*^2^ of ‘naive’ and ‘relaxed’ sparse RRR solutions (Kobak et al., 2019) for various elastic net penalties (*α* and *λ*). ‘Relaxed’ means that the model was re-fit without a lasso penalty using only the selected genes; ‘naive’ means that it was not re-fit. Vertical dashed lines at 25 genes corresponds to the choice made for Figure 3. The best performance is around ~100 genes, but we chose 25 for the sake of interpretability. The subsequent panels only show results for the ‘relaxed’ models. **(b)** Cross-validated *R*^2^ using *α* = 1 for different ranks from rank 1 to rank 16 (full rank). **(c)** Cross-validated *R*^2^ using *α* = 1 and *λ* needed to obtain 25 genes for different ranks. The peak performance is achieved with rank ~12 (inset), but rank-5 model used in the main text is almost as good. **(d)** Cross-validated correlations between sequential projections of the transcriptomic and electrophysiological data sets (rank-5 models with *α* = 1). For any given number of selected genes, correlations decrease monotonically for higher components.

**Figure S9:**
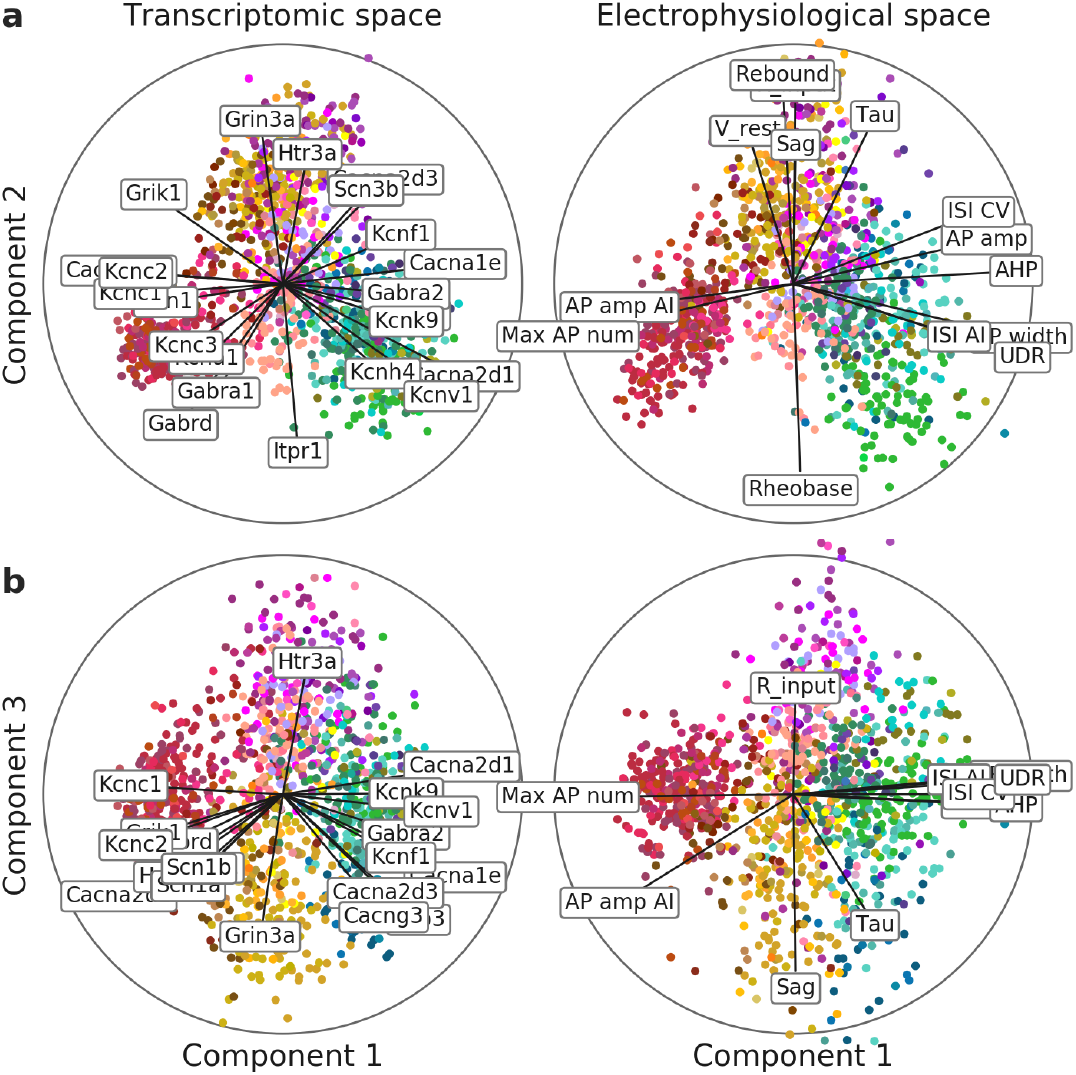
Reduced-rank regression model using only ion channel genes. A full analogue of Figure 3 but using only 328 ion channel genes (https://www.genenames.org/data/genegroup/#!/group/177), of which 293 were detected in our dataset in at least 10 cells.

**Figure S10:**
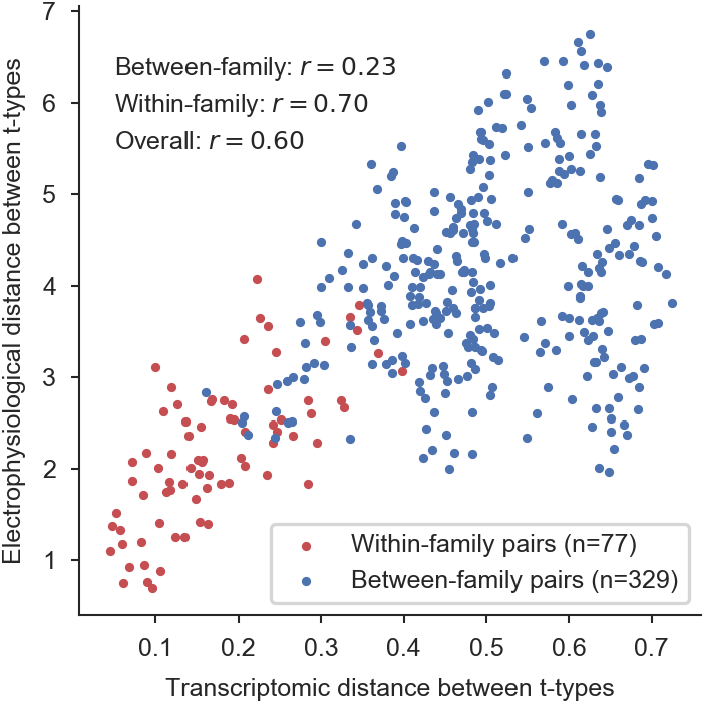
Pooled within-family analysis. The same analysis as in Figure 5a,b (insets) but pooled across all families. We found 29 t-types with at least 10 cells, and computed transcriptomic and electrophysiological distances between all pairs. 77 pairs had t-types belonging to the same family (within-family pairs) and the correlation across those pairs was *r* = 0.70.

**Figure S11:**
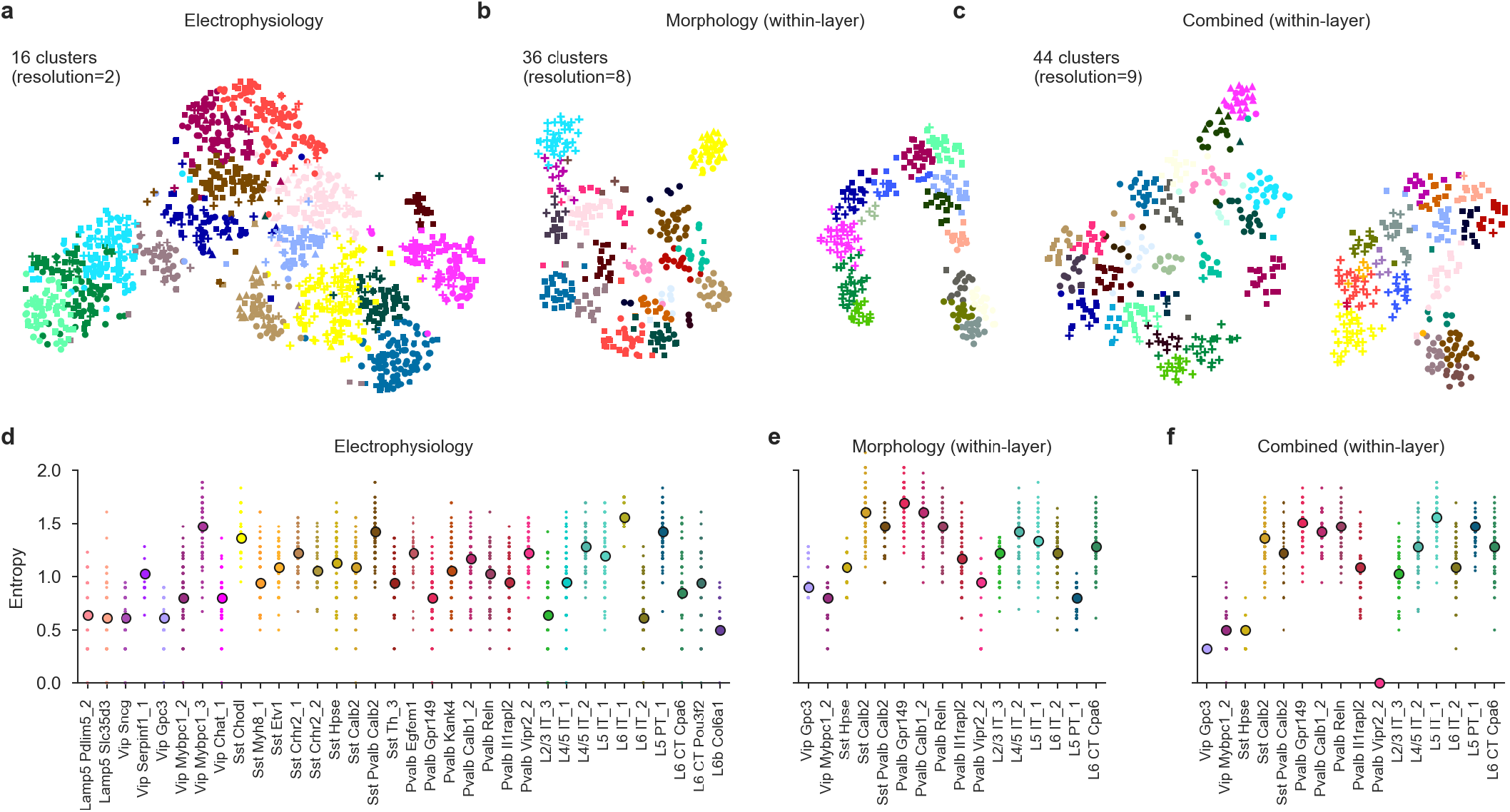
Clustering entropies. **(a–c)** We used Leiden clustering (Traag et al., 2019) to cluster the cells based on electrophysiological, morphological, and combined features. The clustering resolution was adjusted to roughly match the number of e-types, m-types, and em-types from Gouwens et al. (2019). The cluster colors in these panels are arbitrary and not the same as the colors used for t-types. **(d–f)** For each t-type with at least 10 cells, we measured the entropy of the cluster assignments. Entropy zero corresponds to all cells getting into one cluster. Higher entropies mean that cells get distributed across many clusters. We repeated the clustering 100 times with different random seeds, and for each of them, subsampled each t-type to 10 cells to measure the entropy. Points show 100 repetitions, big markers show medians. When using morphological and combined features, all t-types were layer-restricted, as in Figure 6. The t-type colors do not correspond to the colors in panels (a–c).

**Figure S12:**
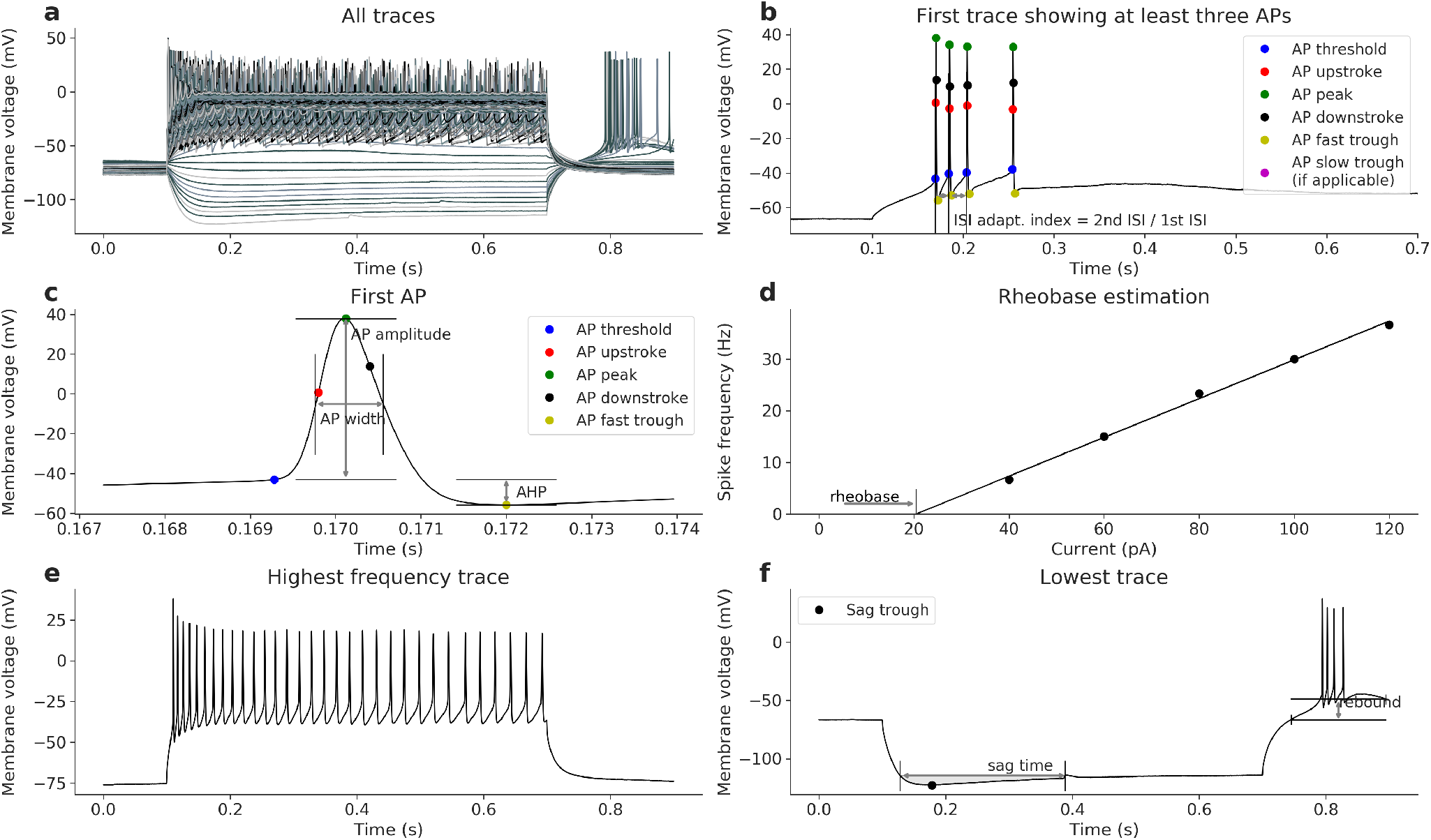
Extraction of the electrophysiological features. All panels show data from the same exemplary cell. **(a)** Membrane potential responses to the consecutive step current injections. Hyperpolarizing currents were used to compute the input resistance (274.80 MOhm) and membrane time constant tau (21.95 ms). **(b)** The first five traces showing spikes were used to compute ISI adaptation index (1.26), ISI average adaptation index (1.15), AP amplitude adaptation index (0.91) and AP amplitude average adapation index (0.99). **(c)** The first AP elicited in this neuron. It was used to compute AP threshold (−40.18 mV), AP amplitude (81.17 mV), AP width (0.80 ms), AHP (−12.60 mV), ADP (0 mV), UDR (1.62) and latency of the first spike (69.28 ms). **(d)** Regression line gives the rheobase estimate (20.44 pA). **(e)** The highest firing trace with 32 APs. This trace was used to estimate the ISI CV (0.27), ISI Fano factor (0.0014 ms), AP CV (0.17) and AP Fano factor (1.32 mV). **(f)** The lowest hyperpolarization trace was used to compute the sag ratio (1.17), sag time (0.26 ms), sag area (31.16 mV·ms) and rebound (17.84 mV).

## Supplementary Tables

**Table S1:**
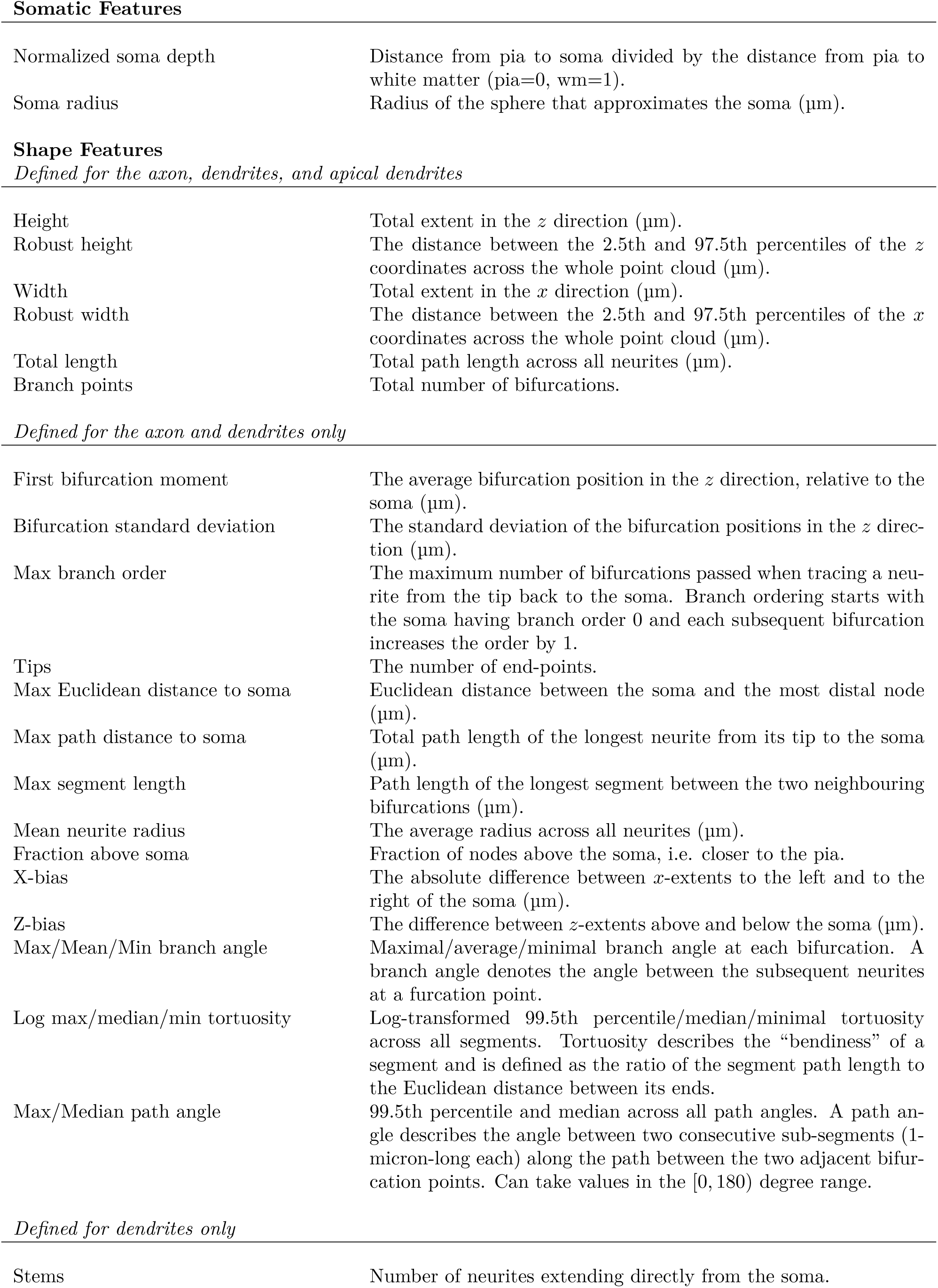
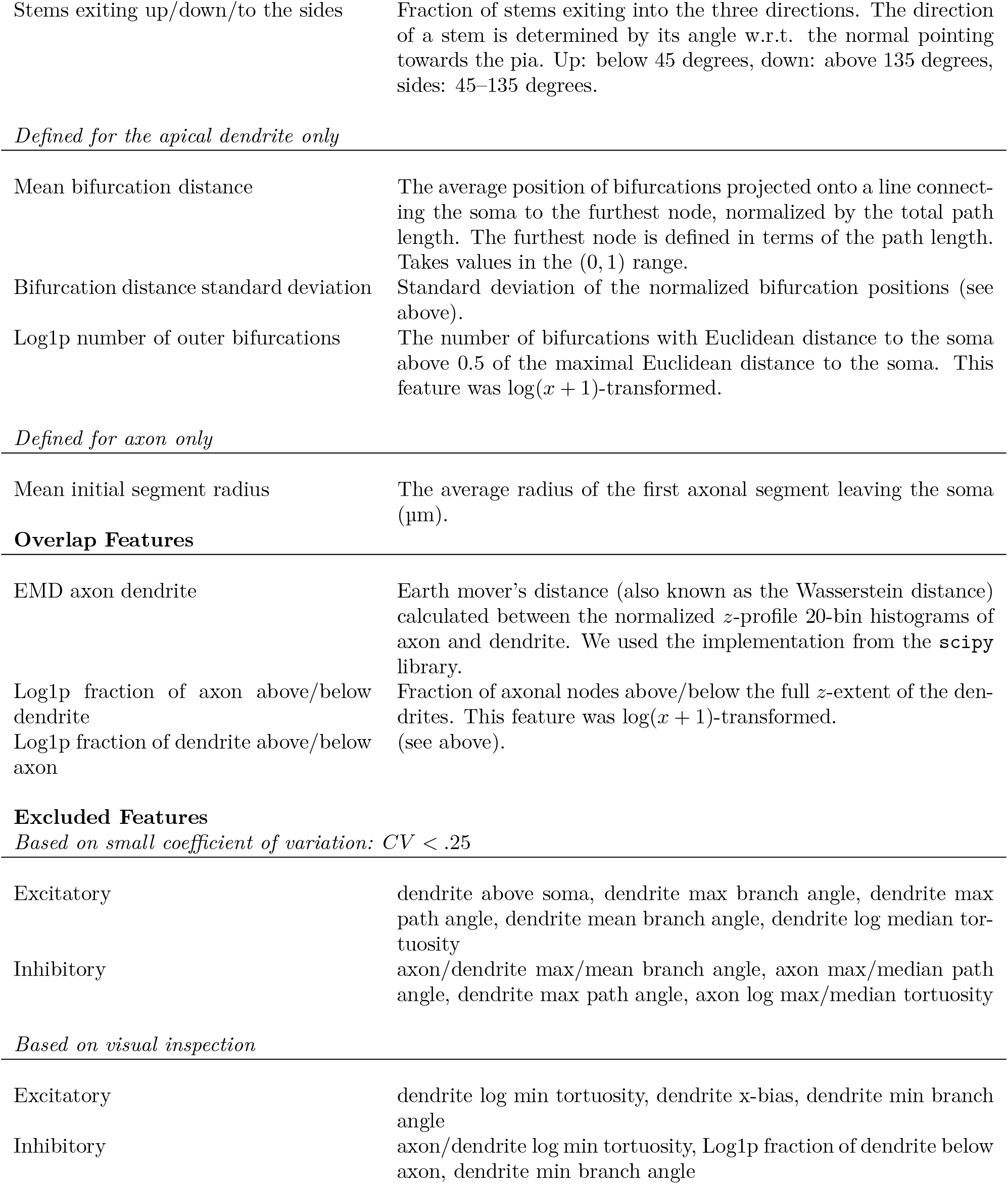
Full list of computed morphometric statistics. It is indicated whether a feature was computed on the soma, the axon, the dendrites or the apical dendrite. The apical dendrite was identified in an automated fashion as the one dendrite with the longest total path length. Overlap features are computed on inhibitory cells only. The section Excluded Features lists all features that have been subsequently excluded.

Somatic Features
|  |  |
| --- | --- |
| Normalized soma depth | Distance from pia to soma divided by the distance from pia to white matter (pia=0, wm=1). |
| Soma radius | Radius of the sphere that approximates the soma ($\mu\text{m}$). |
| Height | Total extent in the $z$ direction ($\mu\text{m}$). |
| Robust height | The distance between the 2.5th and 97.5th percentiles of the $z$ coordinates across the whole point cloud ($\mu\text{m}$). |
| Width | Total extent in the $x$ direction ($\mu\text{m}$). |
| Robust width | The distance between the 2.5th and 97.5th percentiles of the $x$ coordinates across the whole point cloud ($\mu\text{m}$). |
| Total length | Total path length across all neurites ($\mu\text{m}$). |
| Branch points | Total number of bifurcations. |
| *Defined for the axon and dendrites only* |  |
| First bifurcation moment | The average bifurcation position in the $z$ direction, relative to the soma ($\mu\text{m}$). |
| Bifurcation standard deviation | The standard deviation of the bifurcation positions in the $z$ direction ($\mu\text{m}$). |
| Max branch order | The maximum number of bifurcations passed when tracing a neurite from the tip back to the soma. Branch ordering starts with the soma having branch order 0 and each subsequent bifurcation increases the order by 1. |
| Tips | The number of end-points. |
| Max Euclidean distance to soma | Euclidean distance between the soma and the most distal node ($\mu\text{m}$). |
| Max path distance to soma | Total path length of the longest neurite from its tip to the soma ($\mu\text{m}$). |
| Max segment length | Path length of the longest segment between the two neighbouring bifurcations ($\mu\text{m}$). |
| Mean neurite radius | The average radius across all neurites ($\mu\text{m}$). |
| Fraction above soma | Fraction of nodes above the soma, i.e. closer to the pia. |
| X-bias | The absolute difference between $x$-extents to the left and to the right of the soma ($\mu\text{m}$). |
| Z-bias | The difference between $z$-extents above and below the soma ($\mu\text{m}$). |
| Max/Mean/Min branch angle | Maximal/average/minimal branch angle at each bifurcation. A branch angle denotes the angle between the subsequent neurites at a furcation point. |
| Log max/median/min tortuosity | Log-transformed 99.5th percentile/median/minimal tortuosity across all segments. Tortuosity describes the “bendiness” of a segment and is defined as the ratio of the segment path length to the Euclidean distance between its ends. |
| Max/Median path angle | 99.5th percentile and median across all path angles. A path angle describes the angle between two consecutive sub-segments (1-micron-long each) along the path between the two adjacent bifurcation points. Can take values in the [0, 180) degree range. |
| *Defined for dendrites only* |  |
| Stems | Number of neurites extending directly from the soma. |
| Stems exiting up/down/to the sides | Fraction of stems exiting into the three directions. The direction of a stem is determined by its angle w.r.t. the normal pointing towards the pia. Up: below 45 degrees, down: above 135 degrees, sides: 45–135 degrees. |
| *Defined for the apical dendrite only* |  |
| Mean bifurcation distance | The average position of bifurcations projected onto a line connecting the soma to the furthest node, normalized by the total path length. The furthest node is defined in terms of the path length. Takes values in the (0, 1) range. |
| Bifurcation distance standard deviation | Standard deviation of the normalized bifurcation positions (see above). |
| Log1p number of outer bifurcations | The number of bifurcations with Euclidean distance to the soma above 0.5 of the maximal Euclidean distance to the soma. This feature was $\log(x + 1)$-transformed. |
| *Defined for axon only* |  |
| Mean initial segment radius | The average radius of the first axonal segment leaving the soma ($\mu\text{m}$). |
| **Overlap Features** |  |
| EMD axon dendrite | Earth mover's distance (also known as the Wasserstein distance) calculated between the normalized $z$-profile 20-bin histograms of axon and dendrite. We used the implementation from the `scipy` library. |
| Log1p fraction of axon above/below dendrite | Fraction of axonal nodes above/below the full $z$-extent of the dendrites. This feature was $\log(x + 1)$-transformed. |
| Log1p fraction of dendrite above/below axon | (see above). |
| Excitatory | dendrite above soma, dendrite max branch angle, dendrite max path angle, dendrite mean branch angle, dendrite log median tortuosity |
| Inhibitory | axon/dendrite max/mean branch angle, axon max/median path angle, dendrite max path angle, axon log max/median tortuosity |

**Table S1:** Full list of computed morphometric statistics.
|  |  |
| --- | --- |
| Excitatory | dendrite log min tortuosity, dendrite x-bias, dendrite min branch angle |
| Inhibitory | axon/dendrite log min tortuosity, Log1p fraction of dendrite below axon, dendrite min branch angle |

## Supplementary Files

Supplementary File 1: A PDF file containing all reconstructed morphologies and electrophysiological traces in the same format as in Figure 2, sorted by transcriptomic type.

