## Supplementary figures and images for "Phenotypic variation within and across transcriptomic cell types in mouse motor cortex"

### Supplementary File 1

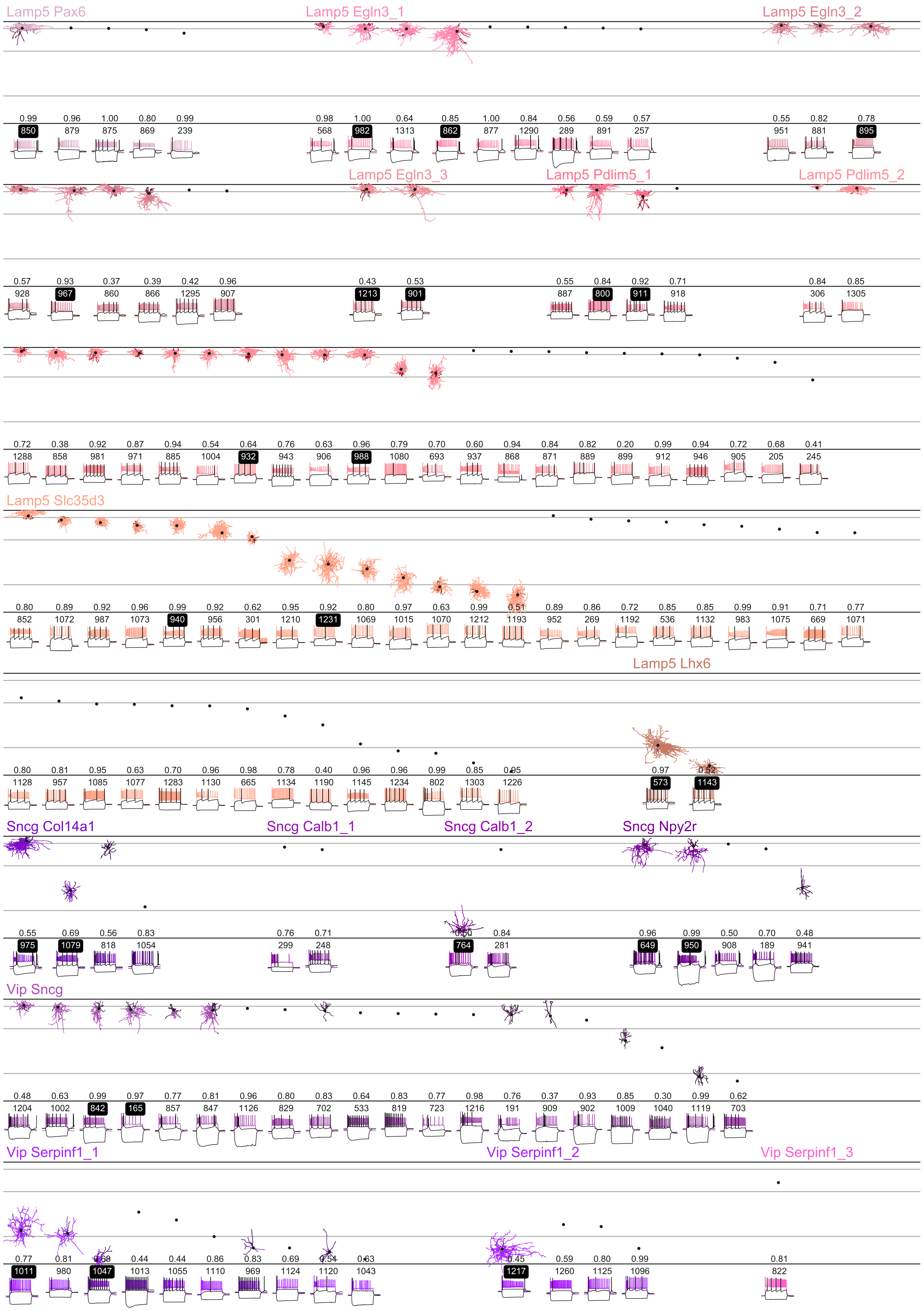

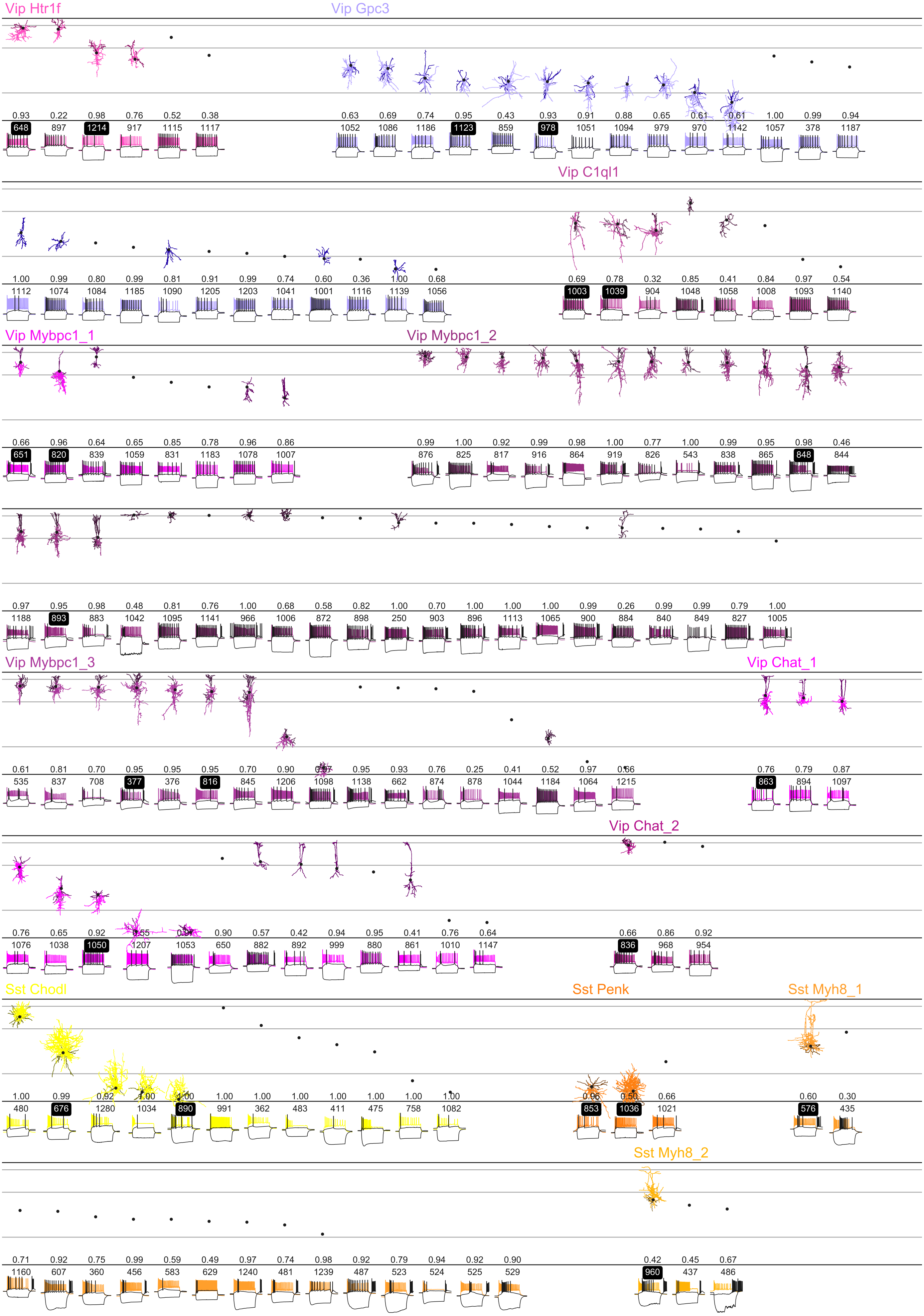

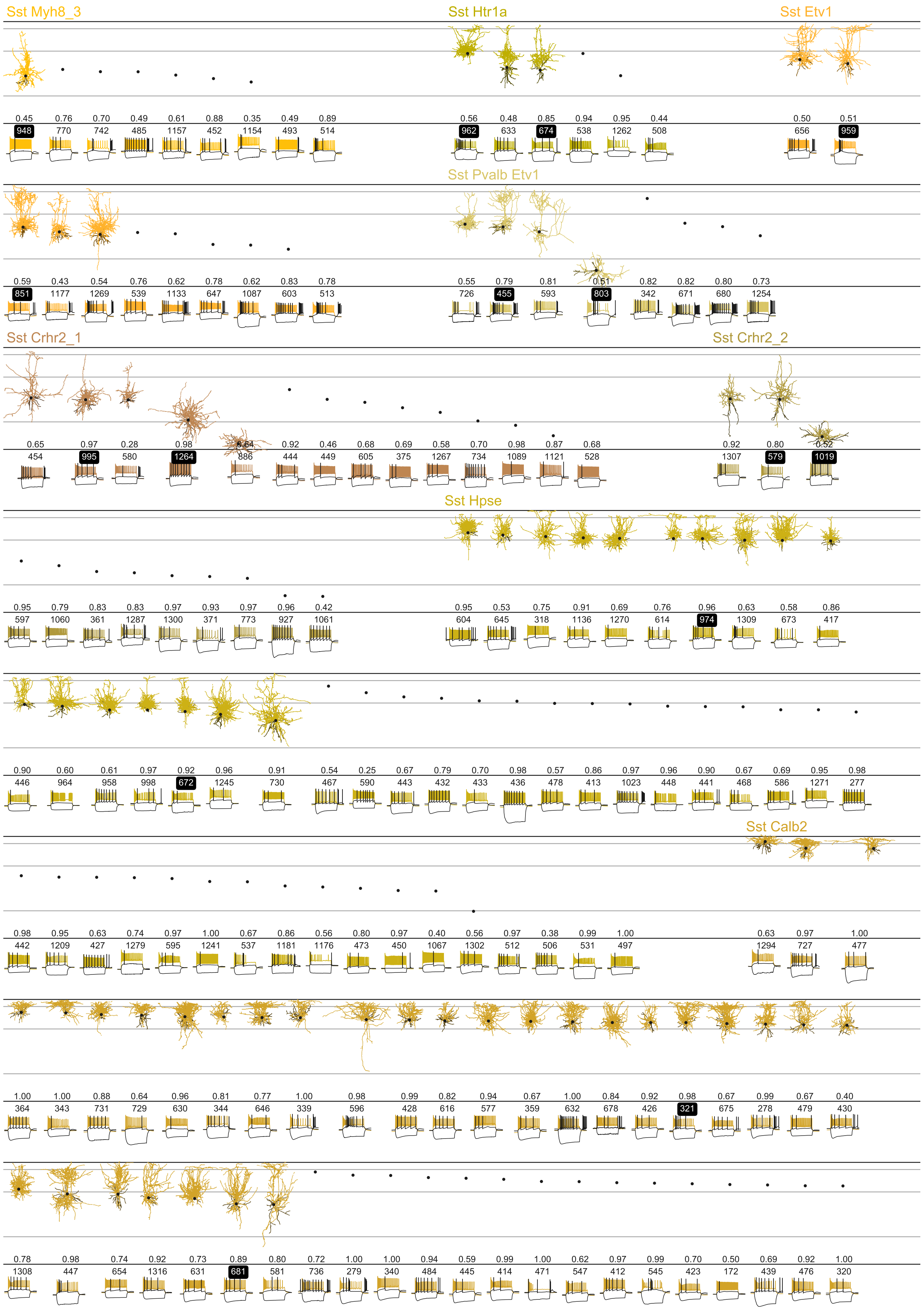

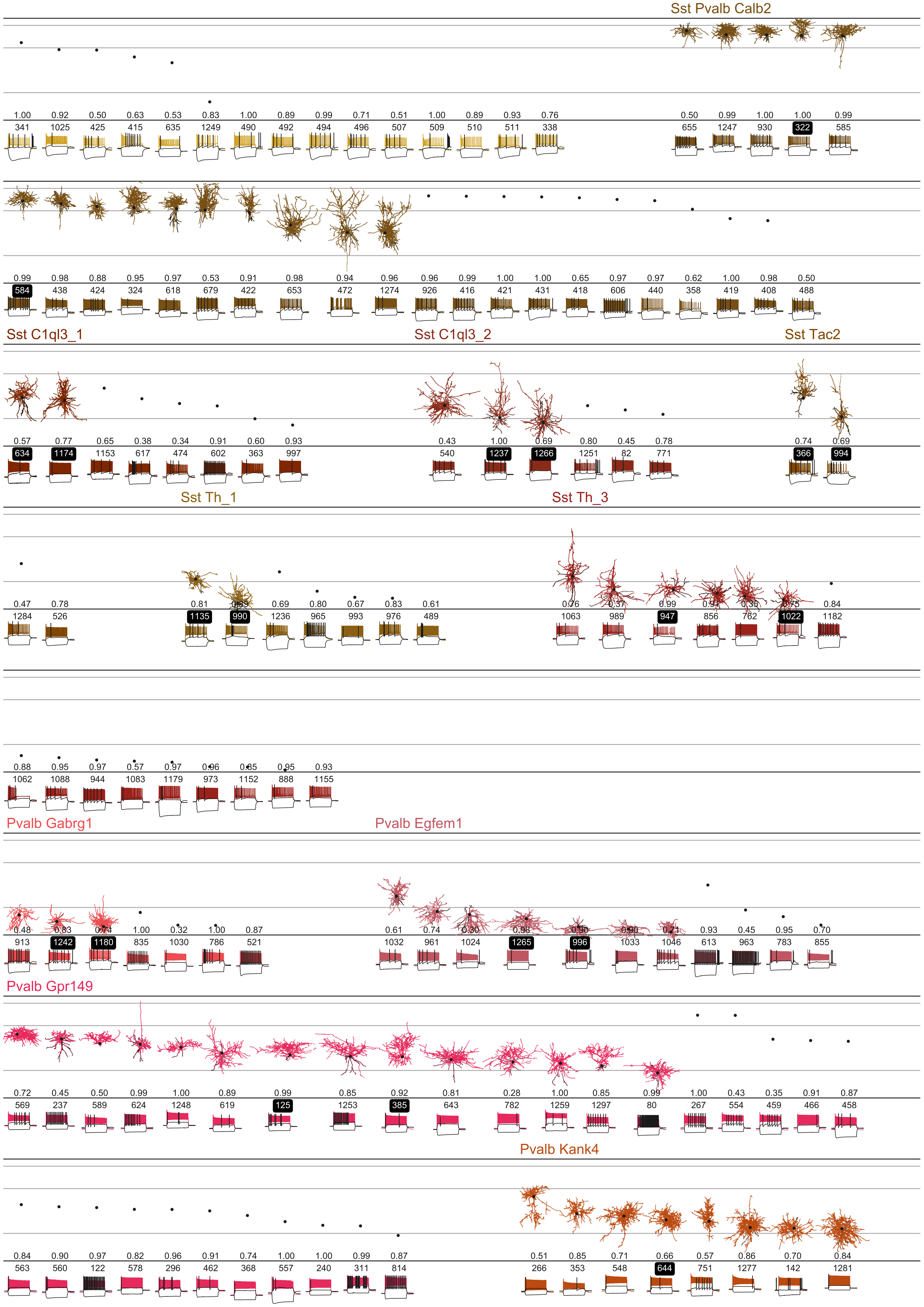

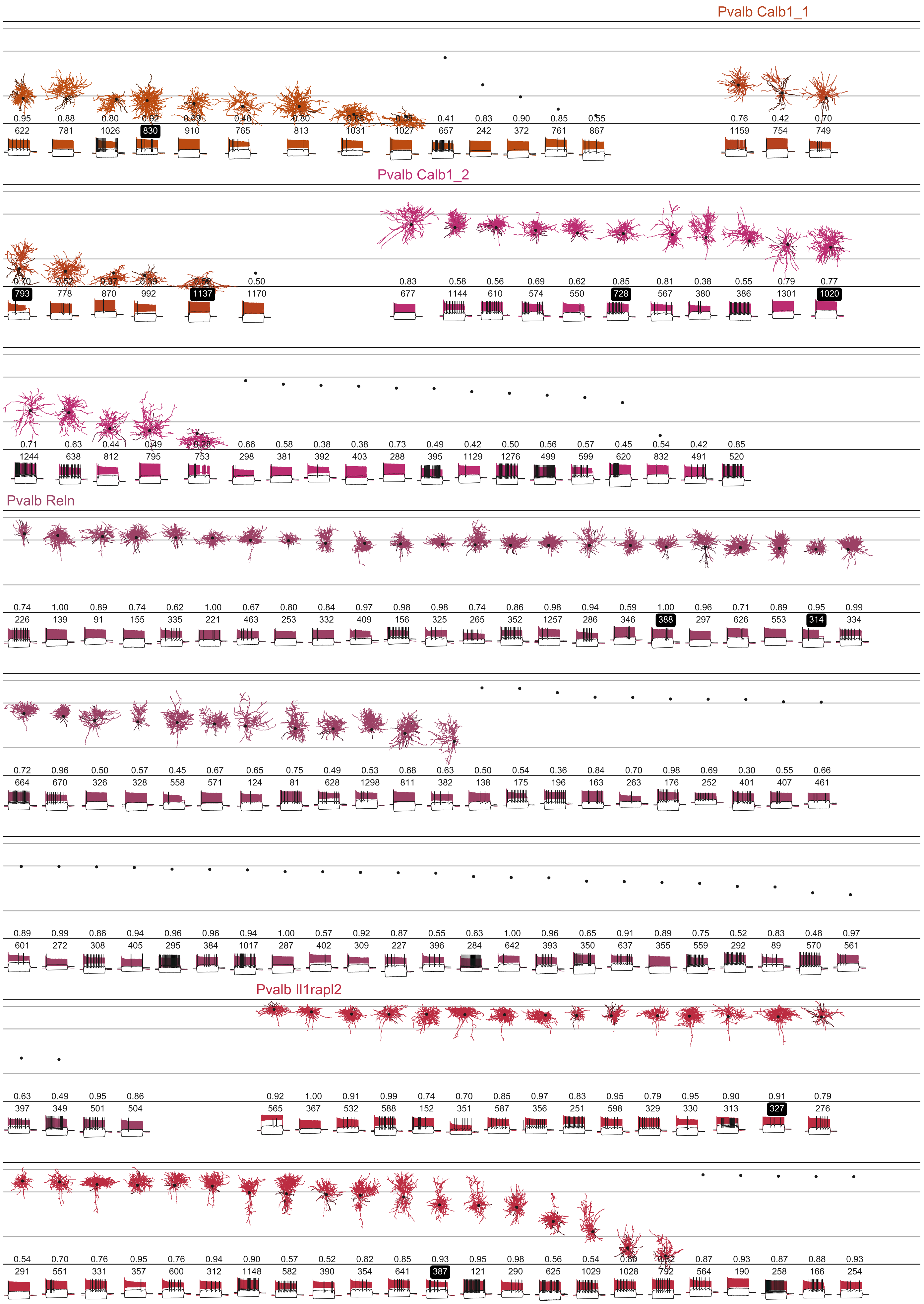

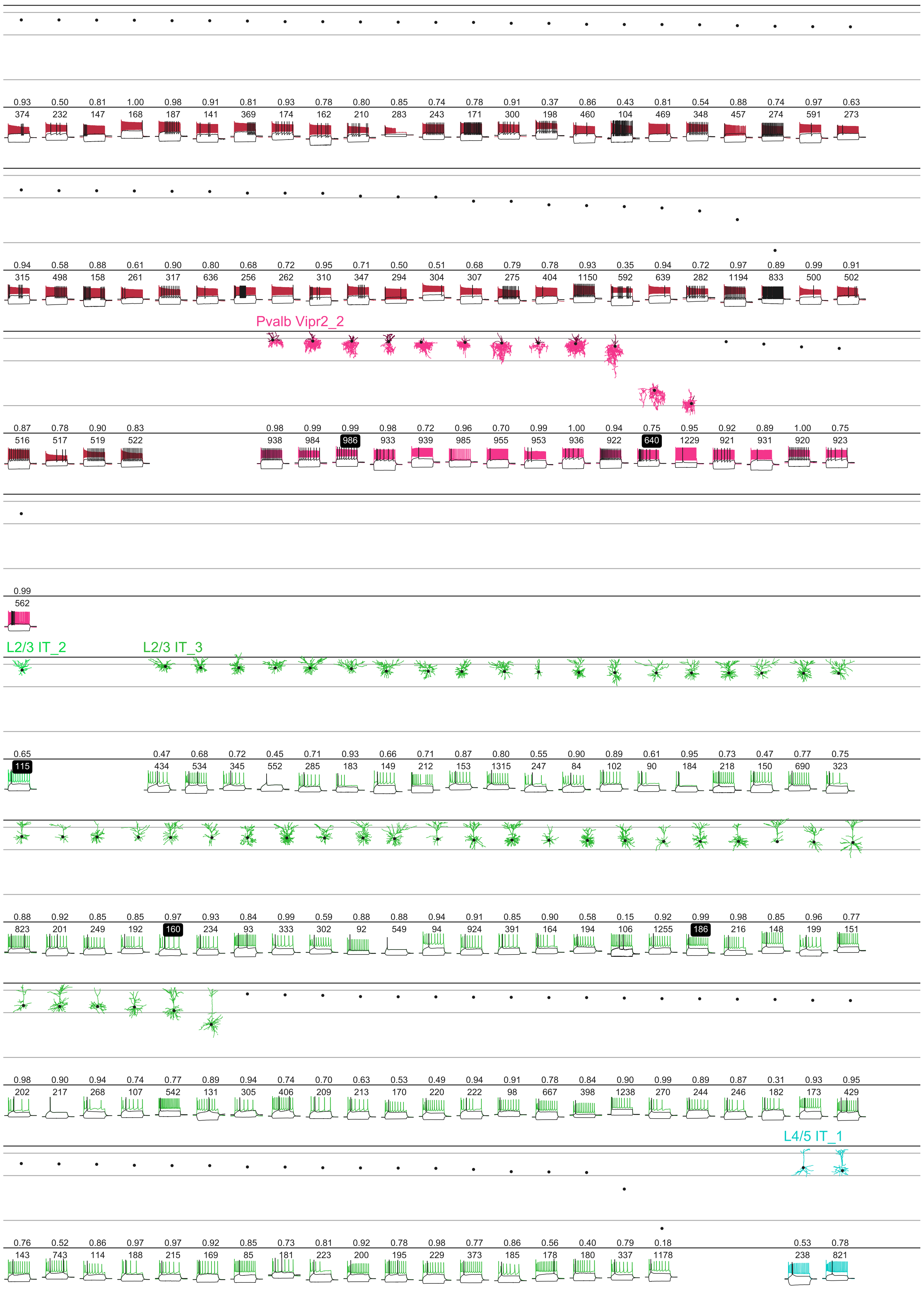

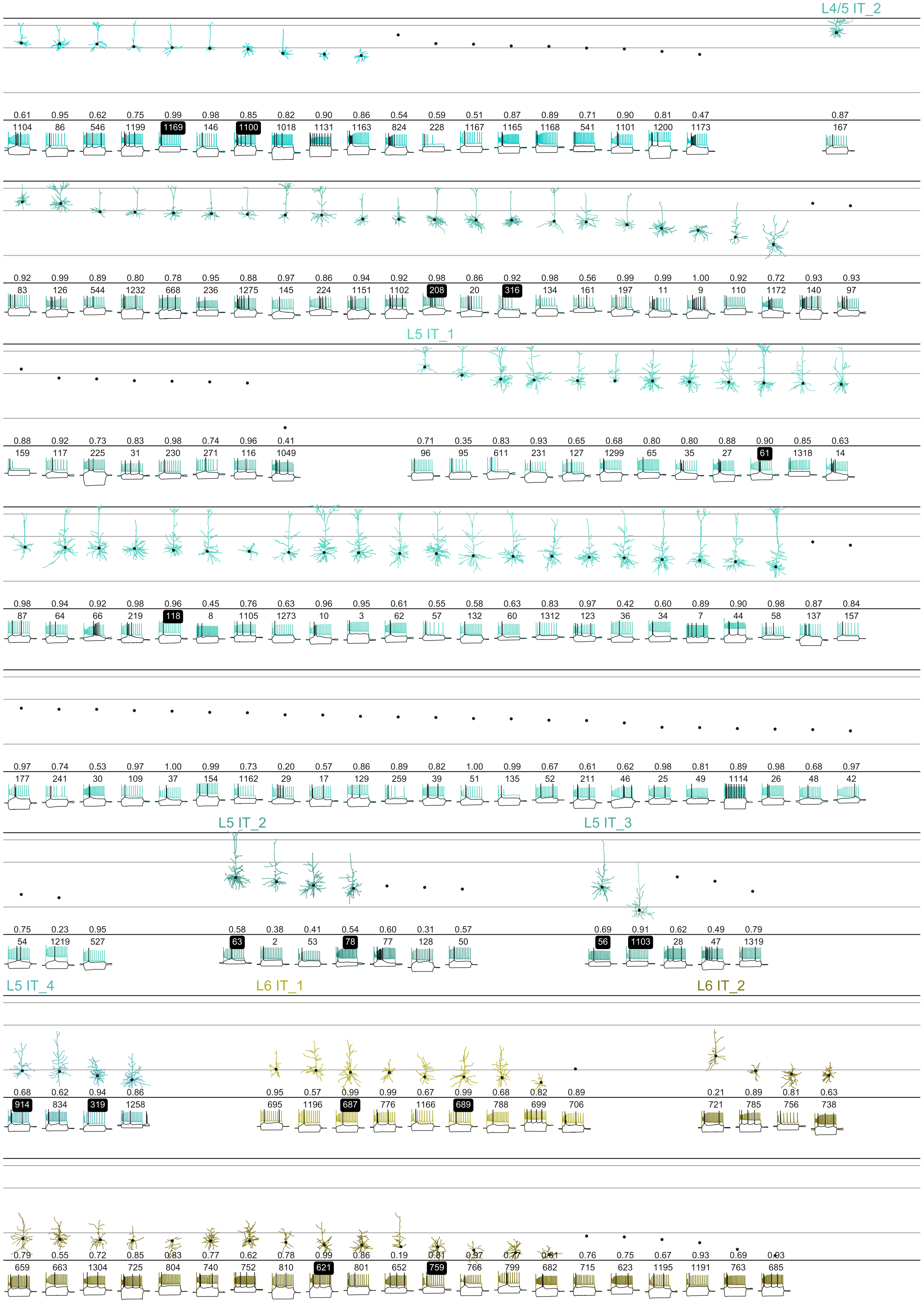

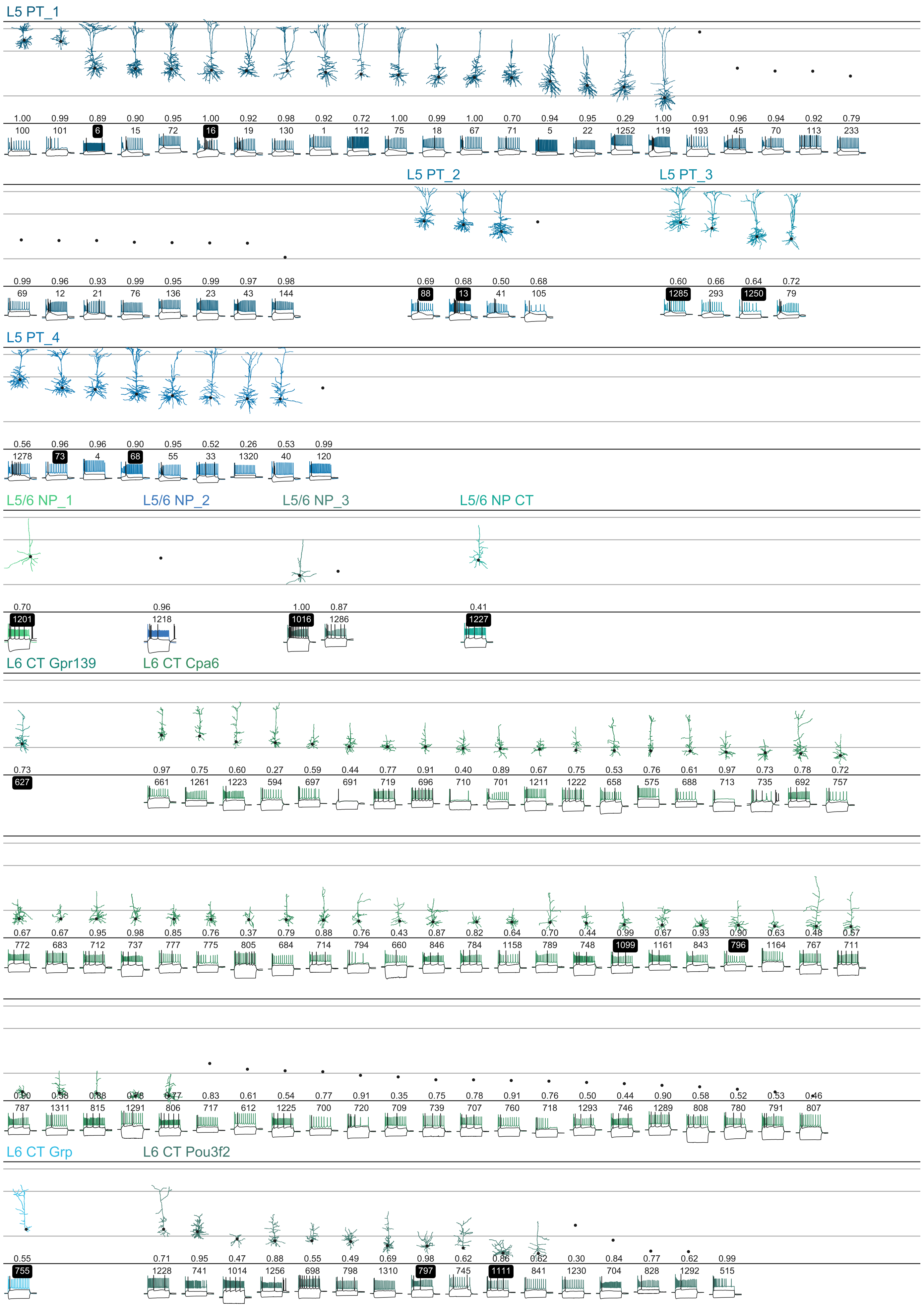

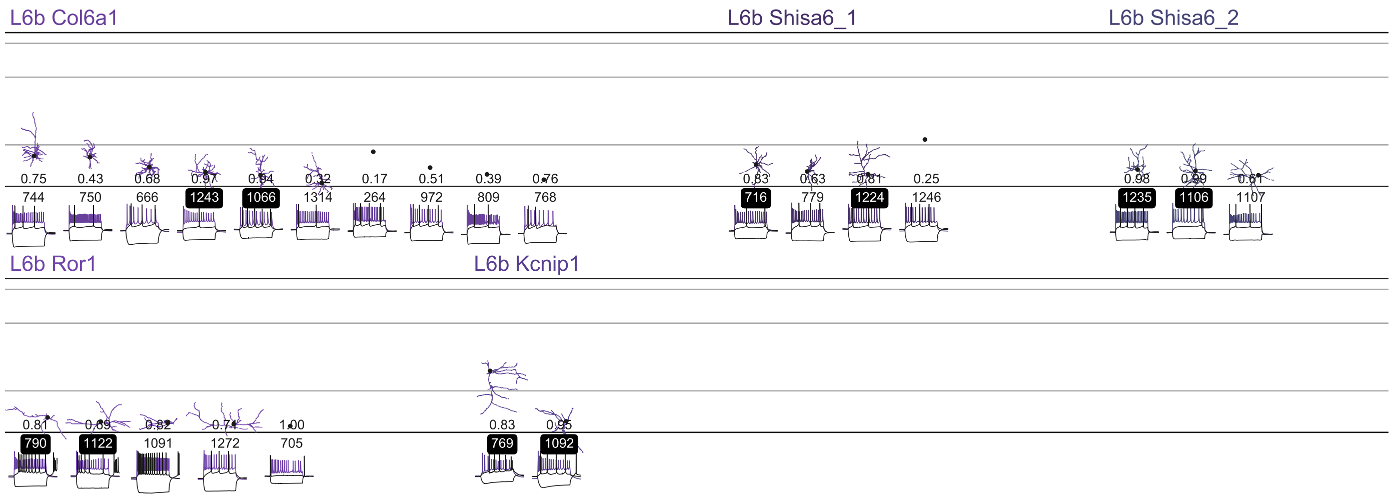
